# Allelic DNA synthesis followed by template switching underlies BRCA1-linked tandem duplication

**DOI:** 10.1101/2024.02.20.581123

**Authors:** Zhi-Cheng Huang, Yi-Li Feng, Qian Liu, Ruo-Dan Chen, Si-Cheng Liu, Meng Wang, An-Yong Xie

## Abstract

Microhomology-mediated short tandem duplication (TD) is among specific mutational signatures associated with *BRCA1*-deficient tumors. Several mechanisms have been proposed for its generation, but may not be applicable in repeat-less regions of the human genome. We thus developed a repeat-less TD reporter and a PCR-based site-specific TD assay to analyze short TDs induced by one-ended DNA double strand breaks (DSBs) converted from DNA nicks in *Brca1*-deficient cells. We found that short TDs induced by DNA nicks are significantly stimulated in *Brca1*-deficient cells. Analysis of TD products revealed that the TD formation is partly mediated by template switching of displaced nascent strand after allelic DNA synthesis. This suggests either allelic DNA synthesis or the strand annealing step of allelic break-induced replication might be more easily aborted in *Brca1-*deficient cells, thus promoting TD. Neither depletion of *Rad51* or *Brca2* nor inactivation of the Brca1 coiled-coil domain stimulated nick-induced TD, indicating that RAD51 loading by BRCA1 is dispensable for BRCA1-mediated TD suppression. These results together provide novel insights into the mechanisms underlying *BRCA1*-linked TD formation in cancer.

## Introduction

Tandem duplication (TD) is considered as an important mechanism for genome evolution and also involved in cancer development (Inaki & Liu, 2012; Alkan *et al*, 2011). Cancer genome sequencing has characterized several classes of cancer-linked TD, one of which is accumulated in *BRCA1*-deficient tumors and has a characteristic span size of less than 50 kb (median value of 11 kb) (Menghi *et al*, 2018, 2016; Nik-Zainal *et al*, 2016; Li *et al*, 2020). Microhomology (MH) is frequently found at the junctions of this TD type and thought to direct the TD formation. Murine mammary cancers deficient for *Brca1* also exhibit similarly short TD that spans 2.5-11 kb (median value of 6.3 kb) (Menghi *et al*, 2018). Like several other mutational signatures and structural variations in *BRCA1*-deficient cancers, this type of TDs could be traced back to defects in the major DNA double strand break repair pathway homologous recombination (HR) and aberrant replication restart at stalled forks, in which BRCA1 plays a key role in end resection, presynaptic RAD51 filament assembly and stabilization of replication fork (Willis *et al*, 2017; Feng *et al*, 2022; Scully *et al*, 2019; Chen *et al*, 2018). This type of TDs has also been exploited as a therapeutic biomarker for the sensitivity to platinum-based chemotherapy and PARP inhibitors in treatment of cancers with HR deficiency (Davies *et al*, 2017; Telli *et al*, 2016).

After identification of TDs associated with *BRCA1* deficiency in cancer genomes, several mechanisms as well as the types of TD-inducing DNA lesions have been proposed and tested in order to explain how short TDs accumulate in *BRCA1*-deficient tumors (Willis *et al*, 2017; Feng *et al*, 2022; Kamp *et al*, 2020). Upon site-specific two-ended DSBs in an HR reporter (i.e., the SCR-RFP reporter) containing a neighboring green fluorescent protein (*GFP*) repeat that serve as a homologous template for non-allelic strand invasion, HR between sister chromatids in combination with long-tract gene conversion (LTGC) generates short TDs, but this TD formation is not elevated by *BRCA1* deficiency (Feng *et al*, 2022; Chandramouly *et al*, 2013; Willis *et al*, 2014). Using the *Escherichia coli* Tus/Ter complex to stall replication forks operating in the 6x*Ter* HR reporter in mammalian cells, Willis and colleagues provided evidence to support that *BRCA1*-linked TDs arise specifically at stalled forks likely *via* a replication restart-bypass mechanism terminated by end joining or by MH-mediated template switching (Willis *et al*, 2017). By inducing site-specific DNA nicks in the original SCR-RFP reporter, we demonstrated that these nicks are converted into one-ended DSBs by DNA replication and generate *BRCA1*-linked mutational signatures including short TDs (Feng *et al*, 2022). Thus, as the most frequent among endogenous DNA lesions (Caldecott, 2022), DNA nicks could be a major source of DNA lesions causing short TDs in *BRCA1*-deficient tumors. While MH-mediated strand invasion is thought to mediate the formation of some TDs induced by DNA nicks in *BRCA1*-deficient cells, it also appears that TD could be induced by template switching or end joining of displaced nascent strand in place of synthesis-dependent strand annealing (SDSA) after allelic strand invasion and allelic DNA synthesis (Feng *et al*, 2022). The end joining could be mediated by DNA polymerase θ (Kamp *et al*, 2020). However, due to the presence of tandem *GFP* repeats in either the 6x*Ter* HR reporter or the SCR-RFP reporter in mammalian cells, the mechanisms proposed thus far for the TD formation may be restricted within at most a half of the human or mouse genome that contains duplicated segments and repetitive elements (Lander *et al*, 2001; Venter *et al*, 2001; Mouse Genome Sequencing Consortium *et al*, 2002) and not be applicable to the other half of the genome lacking repetitive sequences.

When DNA nicks are converted into one-ended DSBs upon collision with DNA replication forks, HR repair of such DSBs is the most likely directed by allelic break-induced replication (BIR), which uses the allele on the sister chromatid as a homologous template for allelic DNA synthesis and is thereby highly faithful in repair (Liu & Malkova, 2022; Anand *et al*, 2013; Llorente *et al*, 2008; Wu & Malkova, 2021). One-ended DSBs may also directly engage MH-mediated BIR (MMBIR), which requires only MH or limited homology to initiate non-allelic strand invasion and proceed with DNA synthesis (Liu & Malkova, 2022; Hastings *et al*, 2009; Sakofsky *et al*, 2015). Due to the identical nature between allelic BIR (aBIR) product and the undamaged allelic substrate in sister chromatids, it is difficult to distinguish the aBIR product from the substrate. However, any interruption in steps of aBIR including end resection, RAD51 loading, allelic strand invasion, allelic D-loop DNA synthesis, displacement of nascent strand, strand annealing or merging of extended D-loop with a converging fork could lead to premature termination of this HR pathway and is expected to have significant impact on repair outcomes of one-ended DSBs converted from DNA nicks. Indeed, previous studies have hinted that enrichment of *BRCA1*-linked TDs could be a consequence of defective BRCA1 function in end resection and strand invasion (Scully *et al*, 2019; Kamp *et al*, 2020). Template switching of displaced nascent strand into non-allelic homology after displacement of nascent strand was also captured in aberrant aBIR repair of one-ended DSBs converted from replication-coupled DNA nicks, generating TD products in *Brca1*-deficient cells (Feng *et al*, 2022). Thus, examining premature termination of aBIR may provide an opportunity to study regulation of aBIR in mammalian cells. It is also unclear whether aBIR is dysregulated in repair of one-ended DSBs converted from DNA nicks in the genome lacking neighboring homologous repeats in *BRCA1*-deficient cells, leading to TD formation, and how.

The TD mechanisms described above were characterized at reporters containing homologous repeats (Willis *et al*, 2017; Feng *et al*, 2022), mimicking repeat-rich genomic sites, and therefore do not represent those at repeat-less regions which comprise approximately 50% of the human genome. Thus, the mechanisms underlying the formation of *BRCA1*-linked TD remains poorly understood in the genome that lacks neighboring repeats. Here, we generated a red fluorescent protein (*RFP*)-based TD reporter measuring the TD formation with no interference of a neighboring homologous repeat in mouse embryonic stem (ES) cells and found significant enrichment of short TDs induced by DNA nicks in *Brca1*-deficient mouse ES cells. We also developed a TD assay at natural genomic sites lacking neighboring homology and found that the level of short TDs induced by DNA nicks was significantly elevated in *Brca1*-deficient cells. Analysis of TD products revealed that the TD formation induced by DNA nicks was mediated by two major mechanisms: 1) MMBIR; 2) aBIR prematurely terminated by template switching-directed MMBIR, abbreviated to aBIR-MMBIR. The latter mechanism suggests that BRCA1 ensure completion of aBIR, thus suppressing TDs. Allelic DNA synthesis in this mechanism shifts the repair junction away from a breakpoint and renders the breakpoint of the genome scarless, indicating that breakpoints may not be equal to junctions characterized with the presence of sequence alterations in genome sequencing, as opposed to the conventional “junctions = breakpoints” rule (Conrad *et al*, 2010; Lam *et al*, 2010; Malhotra *et al*, 2013; Quinlan *et al*, 2010). Further, we demonstrated that RAD51 loading by BRCA1 was not required for TD suppression, suggesting that some *BRCA1*-deficient tumors may not harbor *BRCA1*-linked TD. Together, these results provide a better understanding of the mechanisms underlying TD formation associated with *BRCA1* deficiency in cancer.

## Results

### Template switching associated with aBIR promotes formation of short TDs in a repeat-less TD reporter

Previously, we reported that TD could be generated through two-round strand invasions at the SCR-RFP reporter, the first into allelic *GFP* repeat and the second into non-allelic *GFP* repeat, upon one-ended DSBs converted from DNA nicks by DNA replication as well as upon two-ended DSBs (Feng *et al*, 2022). Given lack of neighboring homologous repeats in majority of a mammalian genome, the use of the SCR-RFP reporter containing two neighboring *GFP* copies (i.e., *TrGFP* and *I-SceI-GFP*) may be unsuitable to mimic TD formation in such part of the genome. We thus used paired sgRNAs with *Streptococcus pyogenes* Cas9 (*Sp*Cas9, abbreviated as Cas9 hereafter) as described previously (Guo *et al*, 2018) to delete *TrGFP* of the SCR-RFP reporter integrated in a single copy at the *Rosa26* locus of mouse ES cells and directly generated a repeat-less TD reporter in the cells (Fig. EV1A).

On the basis of previous finding that *BRCA1*-linked TDs induced by one-ended DSBs are generated by the mechanisms directed by either MH-mediated one-round strand invasion (MH-ORSI) or template switching-associated two-round strand invasion (TRSI) in the SCR-RFP reporter (Feng *et al*, 2022), we proposed that similar mechanisms could be applied to TD induction in the TD reporter upon two-ended DSBs induced by Cas9 or one-ended DSBs induced by Cas9n (Fig. 1A). The cassette including the exon B and A of *RFP* and *I-SceI-GFP* could be duplicated, generating a new cassette with correctly orientated *RFP* exons (i.e., A to B) and making cells *RFP^+^* (Fig. 1A); thus, the frequency of *RFP^+^* cells is used to reflect the level of TD in the repeat-less genomic regions. As replication-coupled one-ended DSBs are primarily repaired by BIR (Liu & Malkova, 2022; Anand *et al*, 2013; Llorente *et al*, 2008; Costantino *et al*, 2014), the signatures of MH-ORSI and template switching-associated TRSI in the HR products indicate the engagement of MMBIR and aBIR followed by template switching in repair of one-ended DSBs for TD formation, respectively (Feng *et al*, 2022). In MMBIR, after one-ended DSBs are generated by collision of nicks with either the leading strand or the lagging strand of a rightward replication fork, the single DNA ends undergo end resection and invade a neighboring region of sister chromatid by reverted MH pairing, directly inducing *RFP^+^* TD events with the predicted span size of 3.4-5.5 kb (Fig. 1A). In aBIR followed by template switching, the resected ends of one-ended DSBs converted from nicks by collision of a rightward replication fork could first allelically invade sister chromatid to start D-loop formation and allelic DNA synthesis. Only after displacement of nascent strands occurs, displaced nascent strands could backtrack and invade a neighboring region of sister chromatid *via* MH-directed template switching as second-round strand invasion, leading to MH-triggered DNA synthesis (i.e., MMBIR) and eventually *RFP^+^* TD with the predicted span size of about 6.9 kb (Fig. 1A). This combined pathway of aBIR and MMBIR is thus termed as the aBIR-MMBIR model (Fig. 1A).

**Figure 1.**
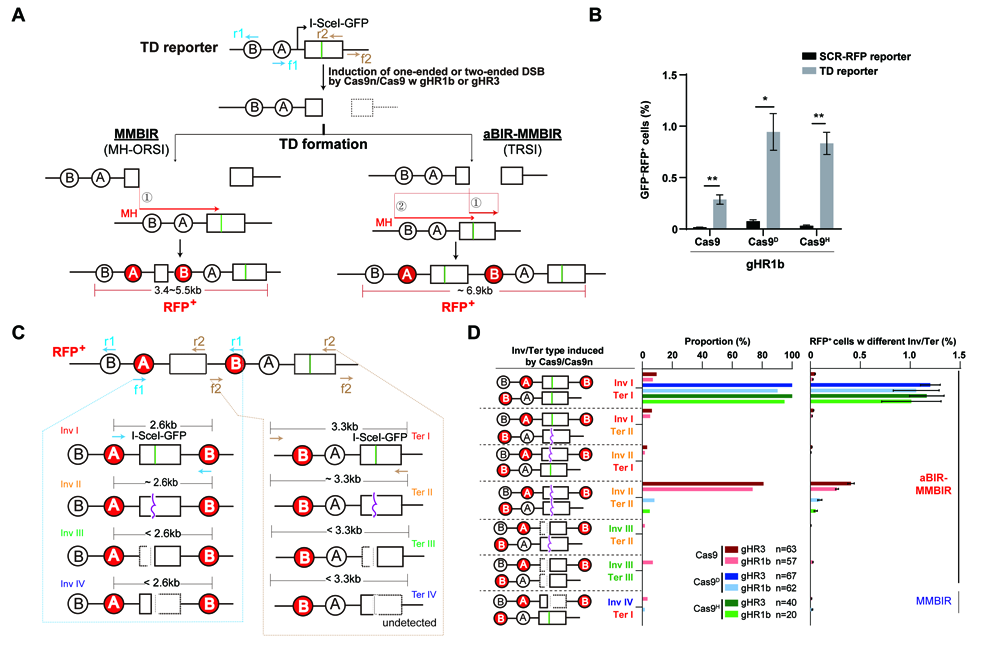
TD induced by Cas9 and Cas9n contains allelic DNA synthesis in the TD reporter in mouse ES cells. **(A)** Two candidate mechanisms underlying Cas9-and Cas9n-induced TD in the TD reporter. In the MMBIR model, MH-mediated strand invasion of left end into the upstream of the *RFP* exon B allows duplication of the *RFP* exon B and A cassette upon site-specific DSBs induced by Cas9 and Cas9n, generating *RFP^+^* cells. The aBIR-MMBIR model involves two rounds of strand invasion (TRSI), first allelic and second non-allelic into sister chromatid to induce TD, some of which generates *RFP^+^* cells. Thus, as a TD outcome, induced *RFP^+^* cells could be used to indicate TD. Two outward primer pairs f1/r1 and f2/r2 on the TD reporter are indicated by blue arrows and brown arrows, respectively. **(B)** Comparison between the parental SCR-RFP reporter and the TD reporter in mouse ES cells in the frequencies of *RFP^+^* cells induced by Cas9, Cas9^D^ or Cas9^H^ with gHR1b. **(C)** Structural classification of *RFP^+^* TD products. Assuming that TD-inducing strand invasion started with left end of Cas9-or Cas9n-induced DSBs, three types of allelic strand invasion junctions, i.e., Inv I, Inv II and Inv III, and one MH-mediated strand invasion junctions, i.e., Inv IV, can be determined and classified by PCR with the outward primer pair f1/r1 as indicated by blue arrows, and four types of termination junctions, i.e., Ter I, Ter II, Ter III and Ter IV, with the outward primer pair f2/r2 as indicated by brown arrows. The straight green line and curve purple line in *I-SceI-GFP* indicate an intact and mutated I-SceI site, respectively. The dotted square in Inv III, Inv IV, Ter III and Ter IV denotes deletion. **(D)** Distribution of Cas9-and Cas9n-induced *RFP^+^* clones with each Inv/Ter type. On the left is schematics of Inv/Ter types detected in *RFP^+^* cells, in the middle the proportion of each Inv/Ter type in *RFP^+^* cells and on the right the frequency of *RFP^+^* cells with each Inv/Ter type. TDs with Inv I/Ter I, Inv I/Ter II, Inv II/Ter I, Inv II/Ter II, Inv III/Ter II and Inv III/Ter III are classified as products of aBIR-MMBIR because nascent DNA from allelic DNA synthesis was found in these TDs. *RFP^+^* cells with Inv IV/Ter I could be induced by MH-mediated one-round strand invasion and are therefore classified as a product of MMBIR. The frequency of *RFP^+^* cells with each Inv/Ter type is calculated as the overall frequency of *RFP^+^*cells × the proportion of the Inv/Ter type in *RFP^+^* cells. Columns indicate the mean ± S.D. from three independent experiments in **B** and **D**. In **D**, n indicates the number of *RFP^+^* clones analyzed. Statistical analysis is performed by paired two-tailed Student’s t test in **B**. *, *P*<0.05; **, *P*<0.01.

To test the functionality of the TD reporter, we used Cas9 or Cas9n (i.e., Cas9^D^ for Cas9 D10A mutant and Cas9^H^ for Cas9 H840A mutant) with the sgRNA gHR1b to induce two-ended or one-ended DSBs at the same target site of *I-SceI-GFP*, respectively. Compared to the negligible level of spontaneous *RFP^+^*cells in TD reporter cells, the frequencies of *RFP^+^* cells induced by Cas9, Cas9^D^ and Cas9^H^ were clearly detectable by flow cytometry (Fig. EV1B). Moreover, these frequencies of *RFP^+^*cells induced by Cas9, Cas9^D^ and Cas9^H^ with gHR1b in the TD reporter were nearly 10-fold higher than those of respective *GFP^−^RFP^+^* cells induced at the same site in parental SCR-RFP reporter mouse ES cells (Fig. 1B). This change confirms that neighboring homologous repeats strongly compete with MH for strand invasion and severely interfere with MH-mediated TD formation.

In the *RFP^+^* TD products derived from the MMBIR model and the aBIR-MMBIR model, the first *GFP* is expected to carry an invasion point and the second *GFP* should harbor the termination point (Fig. 1A). To determine the invasion and termination types of *RFP^+^* cells, we sorted *RFP^+^* cells, picked single *RFP^+^* clones and analyzed the sequences of tandem *GFP* copies in these *RFP^+^* clones after targeted PCR amplification with primer pairs f1/r1 or f2/r2 (Fig. 1A and C). Among 63 *RFP^+^* clones induced by Cas9-gHR3, 57 by Cas9-gHR1b, 67 by Cas9^D^-gHR3, 62 by Cas9^D^-gHR1b, 40 by Cas9^H^-gHR3 and 20 by Cas9^H^-gHR1b (Fig. 1D), we found four types of invasion and termination (Fig. 1C): 1) Type I of invasion (Inv I) or termination (Ter I) with no scars at the breakpoint of the respective *GFP* copy; 2) Type II of invasion (Inv II) or termination (Ter II) with scars (i.e., small indels) at the breakpoint of the respective *GFP* copy; 3) Type III of invasion (Inv III) or termination (Ter III) with deletion of the *GFP* left portion from the breakpoint of the respective *GFP* copy; 4) Type IV of invasion (Inv IV) or termination (Ter IV) with deletion of the *GFP* right portion from the breakpoint of the respective *GFP* copy. Among *RFP^+^* TD products analyzed, aBIR-MMBIR could generate the type with Inv I/Ter I, Inv I/Ter II, Inv II/Ter I, Inv II/Ter II, Inv III/Ter II and Inv III/Ter III whereas Inv IV/Ter I was supposedly induced by MMBIR (Fig. 1A-D).

Cas9-induced *RFP^+^* clones are quite different from Cas9n-induced *RFP^+^* clones as majority of Cas9-induced *RFP^+^* clones were Inv II/Ter II whereas Cas9n-induced *RFP^+^* clones were mostly associated with Inv I/Ter I (Fig. 1D). In particular, for each Cas9-induced *RFP^+^* clone with Inv II/Ter II, both the Inv II point and the Ter II point shared the same indels (Table EV1). Interestingly, an Inv III/Ter III product also contained the same deletion of the *GFP* left portion in the invasion point and the termination point (Table EV 1). The fact that the invasion point and the termination point of *RFP^+^* TD shares the same sequences supports the involvement of allelic DNA synthesis prior to template switching in the formation of TD with the Inv III/Ter III type as with Inv I/Ter I and Inv II/Ter II. As explained previously (Feng *et al*, 2022), the *I-SceI-GFP* copy in the sister chromatid template could first be cleaved by Cas9 and mutated by non-homologous end joining (NHEJ) prior to allelic strand invasion, and the mutation generated could then be introduced into the nested *GFP* in TD products with Inv II/Ter II and Inv III/Ter III by allelic DNA synthesis. Therefore, Inv III/Ter III is considered as a type of Inv II/Ter II. However, Inv III/Ter III was infrequent as were the other combinations of invasion and termination types, such as Inv I/Ter II, Inv II/Ter I, Inv III/Ter II and Inv IV/Ter I (Fig. 1D). In contrast, *RFP^+^* cells with Inv IV/Ter I were the only MMBIR product detected and the frequency of these *RFP^+^*cells was minimal (Fig. 1D), indicating little involvement of MMBIR in the TD formation in the TD reporter. Thus, aBIR-MMBIR appears to be a dominant pathway for the TD formation induced by two-ended DSBs and replication-coupled one-ended DSBs in the TD reporter.

### BRCA1 suppresses TD initiated by template switching-associated TRSI

To further determine the mechanisms underlying BRCA1-mediated suppression of TD formation, we analyzed TDs induced by Cas9, Cas9^D^ and Cas9^H^ together with gHR1b at the TD reporter site in mouse ES cells depleted of *Brca1* (Fig. EV2A and EV2B). *RFP^+^* cells induced by Cas9 and Cas9n were all increased by *Brca1* depletion (Fig. 2B). We then disrupted exon 15 encoding part of the BRCT domain of Brca1 by paired Cas9 approach as previously described (Feng *et al*, 2022; Guo *et al*, 2018) to generate 3 *Brca1*-deficient (*Brca1^m/m^*) TD reporter mouse ES clones as well as 3 isogenic wild-type (*Brca1^+/+^*) control clones (Fig. EV3A). In line with previous reports, lack of the BRCT domain resulted in degradation of BRCA1 protein in *Brca1^m/m^* cells (Fig. EV3B) (Williams *et al*, 2003). By comparing 3 isogenic *Brca1^m/m^* clones with 3 isogenic *Brca1^+/+^* clones, the frequencies of *RFP^+^* cells induced by Cas9, Cas9^D^ and Cas9^H^ with gHR1b were all significantly elevated by *Brca1* deficiency (Fig. EV3C-E), indicating *Brca1*-mediated TD suppression. However, consistent with previous finding (Feng *et al*, 2022), *Brca1* deficiency did not elevate the frequencies of *RFP^+^* cells induced by Cas9 or Cas9^D^ with gHR1b in parental SCR-RFP reporter mouse ES cells (Fig. EV3F), again suggesting the potential difference in regulating TD formation between genomic regions with and without neighboring repeats. We tested gHR3, gHR1b and 7 other sgRNAs targeting different sites in the TD reporter in a *Brca1^+/+^* clone (i.e., #2) and a *Brca1^m/m^* clone (i.e., #113) (Fig. 2A and EV4A) and found that *Brca1* deficiency consistently stimulated the production of *RFP^+^*cells induced by Cas9, Cas9^D^ and Cas9^H^ (Fig. 2B and EV4B-D), further confirming *Brca1*-mediated TD suppression.

**Figure 2.**
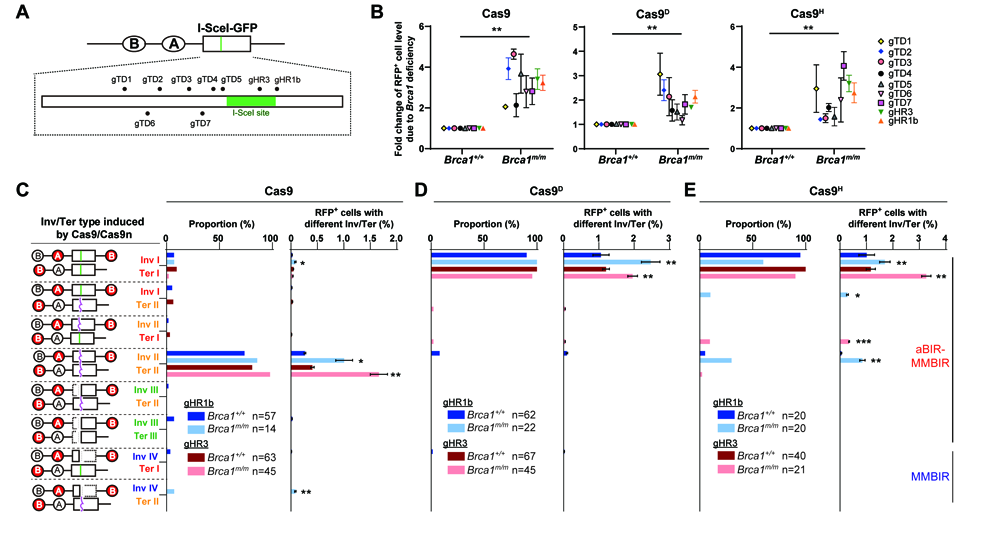
*Brca1* deficiency stimulates Cas9-and Cas9n-induced TD in the TD reporter. **(A)** Schematic of the TD reporter with 9 target sites by Cas9 or Cas9n, with the NGG PAMs for gTD1, gTD2, gTD3, gTD4, gTD5, gHR1b and gHR3 on the Crick strand and for gTD6 and gTD7 on the Watson strand as indicated. **(B)** Fold change in the frequency of the *RFP^+^* cells induced by Cas9 (left), Cas9^D^ (middle) or Cas9^H^ (right) along with indicated sgRNA in *Brca1^+/+^* and *Brca1^m/m^*cells. Fold change of *RFP^+^* cell level due to *Brca1* deficiency is calculated by normalizing the frequencies of Cas9-, Cas9^D^-and Cas9^H^-induced *RFP^+^* cells in *Brca1^+/+^*cells to 1. **(C-E)** Distribution of *RFP^+^* cells with each Inv/Ter type induced by Cas9 (**C**), Cas9^D^ (**D**) and Cas9^H^ (**E**) together with gHR1b and gHR3 in *Brca1^+/+^*and *Brca1^m/m^* cells. The proportion of each Inv/Ter type in *RFP^+^* cells and the frequency of *RFP^+^* cells with each Inv/Ter type are shown. *RFP^+^* cells with Inv I/Ter I, Inv I/Ter II, Inv II/Ter I, Inv II/Ter II, Inv III/Ter II and Inv III/Ter III are classified as products of aBIR-MMBIR, and Inv IV/Ter I and Inv IV/Ter II as MMBIR. The frequency of *RFP^+^* cells with each Inv/Ter type is calculated as the overall frequency of *RFP^+^* cells × the proportion of the Inv/Ter type in *RFP^+^* cells. Each symbol represents the value from at least three independent experiments for indicated sgRNAs in **B**, and statistics is performed by two-tailed Student’s t-test. Columns for the frequency of *RFP^+^* cells with each Inv/Ter type indicate the mean ± S.D. from three or more independent measurements of induced *RFP^+^* cells in **C-E**, and statistics is performed by paired two-tailed Student’s t test. n indicates the number of *RFP^+^* clones analyzed in **C-E**. *, *P*<0.05; **, *P*<0.01; ***, *P*<0.001.

To determine which type of TDs are suppressed by *Brca1*, we first analyzed the tandem *GFP* sequences of 57 *RFP^+^* clones induced by Cas9-gHR1b and 63 *RFP^+^* clones induced by Cas9-gHR3 from *Brca1^+/+^* cells and 14 *RFP^+^* clones induced by Cas9-gHR1b and 45 *RFP^+^* clones induced by Cas9-gHR3 from *Brca1^m/m^* cells (Fig. 2C). TDs were mostly an Inv II/Ter II product, a TD type that is generated by aBIR-MMBIR and also increased by *Brca1* deficiency (Fig. 2C). Thus, the elevation of the Inv II/Ter II products had the most impact on TD enrichment in *Brca1*-deficient cells (Fig. 2C). After examining 62 *RFP^+^* clones induced by Cas9^D^-gHR1b from *Brca1^+/+^* cells, 22 *RFP^+^* clones by Cas9^D^-gHR1b from *Brca1^m/m^* cells, 67 *RFP^+^* clones by Cas9^D^-gHR3 from *Brca1^+/+^* cells, 45 *RFP^+^* clones by Cas9^D^-gHR3 from *Brca1^m/m^* cells, we found that most of TDs were an Inv I/Ter I product induced by aBIR-MMBIR (Fig. 2D). Only this TD type was significantly promoted by *Brca1* deficiency (Fig. 2D), suggesting its dominant contribution to TD formation in *Brca1^m/m^* cells. Analysis of *RFP^+^* clones induced by Cas9^H^ similarly revealed that *Brca1* deficiency mostly stimulated formation of TDs with the Inv I/Ter I type, which is also dominant in *Brca1^+/+^* cells, and the Inv II/Ter II type (Fig. 2E). As both Inv I/Ter I and Inv II/Ter II are generated by aBIR-MMBIR, these results together suggest that *Brca1* deficiency promote template switching-directed MH-mediated second-round strand invasion after nascent DNA strand is displaced during allelic DNA synthesis, leading to break-induced TD formation upon two-ended DSBs or replication-coupled one-ended DSBs.

### Allelic DNA synthesis prior to template switching renders TD-inducing breakpoints scarless

As breakpoints are often defined as junctions characterized with the presence of sequence alterations in genome sequencing (Conrad *et al*, 2010; Lam *et al*, 2010; Malhotra *et al*, 2013; Quinlan *et al*, 2010), we asked whether TD formation initiated at different breakpoints were associated with different junctions in this TD reporter in both *Brca1^+/+^*cells and *Brca1^m/m^* cells. Surprisingly, analysis of TD products revealed that 9.52% (6 out of 63) *RFP^+^* TDs induced by the Cas9-gHR3 breakpoint shared the same Inv I/Ter I TD product with 7.01% (4 out of 57) *RFP^+^* TDs induced by the Cas9-gHR1b breakpoint in *Brca1^+/+^*cells, and 2.22% (1 out of 45) *RFP^+^* TDs induced by the Cas9-gHR3 breakpoint shared the same Inv I/Ter I TD product with 7.14% (1 out of 14) *RFP^+^* TDs induced by the Cas9-gHR1b breakpoint in *Brca1^m/m^*cells (Fig. 3A). Further, 100% (67 out of 67) *RFP^+^* TDs induced by the Cas9^D^-gHR3 breakpoint had the same Inv I/Ter I TD product with 90.32% (56 out of 62) *RFP^+^* TDs induced by the Cas9^D^-gHR1b breakpoint in *Brca1^+/+^* cells, and 95.55% (43 out of 45) *RFP^+^*TDs induced by the Cas9^D^-gHR3 breakpoint shared the same Inv I/Ter I TD product with 100% (22 out of 22) *RFP^+^* TDs induced by the Cas9^D^-gHR1b breakpoint in *Brca1^m/m^* cells (Fig. 3A). Similarly, both the Cas9^H^-gHR3 breakpoint and the Cas9^H^-gHR1b breakpoint generated a high level of the same Inv I/Ter I TD product in both *Brca1^+/+^*cells and *Brca1^m/m^* cells (Fig. 3A). These Inv I/Ter I TD products contained the same MH junction “TGTC” and a scarless breakpoint (Fig. 3A). This indicates that in the Inv I/Ter I product, whose formation is mediated by aBIR-MMBIR, DNA synthesis following allelic strand invasion restore the sequence of the breakpoints by Cas9 or Cas9n and shift the MH-mediated junctions away from the breakpoints. As a result, the breakpoints in these TD products would be impossible to be identified if they were not known as Cas9 or Cas9n target sites.

**Figure 3.**
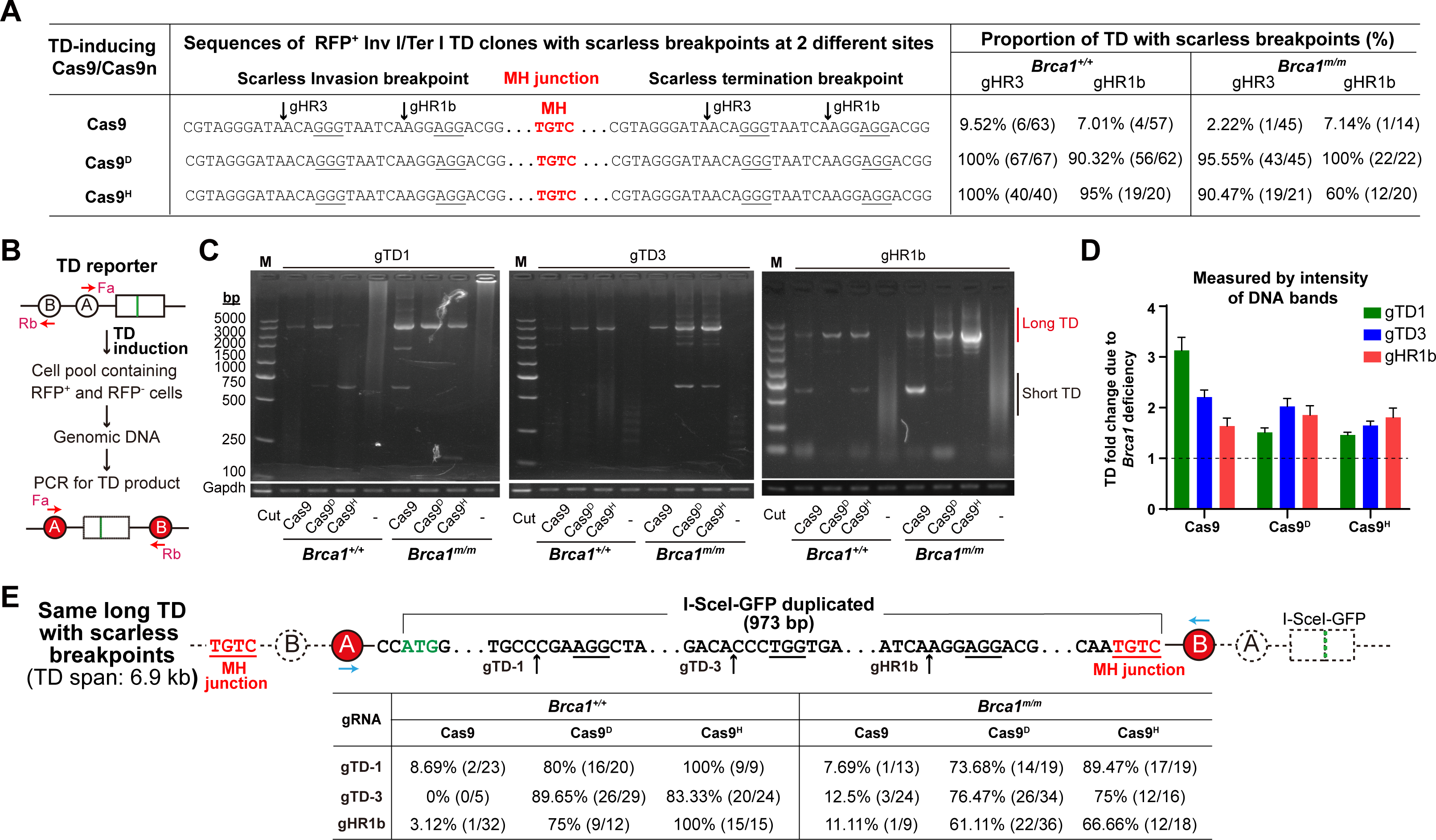
Allelic DNA synthesis in aBIR-MMBIR masks the TD breakpoints in both *Brca1^+/+^* and *Brca1^m/m^*cells. **(A)** Representative sequences and proportions of Cas9-and Cas9n-induced Inv I/Ter I TDs with scarless breakpoints in the TD reporter in *Brca1^+/+^*and *Brca1^m/m^* cells. Breakpoints by Cas9 and Cas9n together with gHR3 and gHR1b are indicated along with MH that mediates TD. The number of TDs with scarless breakpoints and the number of total TD events are shown in parenthesis after each proportion value in percentage. **(B)** Schematic of PCR-based TD detection in the TD reporter. After TD induction by Cas9 or Cas9n in the TD reporter, TD products in a pool of *RFP^+^*cells and *RFP^−^* cells as shown can be amplified by PCR with Fa/Rb, an outward primer pair indicated in red arrows in the TD reporter. **(C)** Representative images of PCR products separated by DNA agarose gel electrophoresis for TDs induced by Cas9, Cas9^D^ or Cas9^H^ together with gTD1 (left), gTD3 (middle) or gHR1b (right) in *Brca1^+/+^*and *Brca1^m/m^* TD reporter cells. PCR products of a *Gapdh* region is used as internal PCR control. Long and short TD products are indicated by red and black vertical lines, respectively. **(D)** Fold change in stimulation of long and short TDs induced by Cas9, Cas9^D^ or Cas9^H^ together with gTD1, gTD3 or gHR1b due to *Brca1* deficiency. The relative TD intensity is regarded as the ratio of the TD intensity to the *Gapdh* intensity measured by ImageJ software. TD fold change due to *Brca1* deficiency is calculated by normalizing the relative TD intensity in *Brca1^+/+^* cells to 1. **(E)** Schematic sequence of 973-bp *I-SceI-GFP* duplicated (top) in the same long TD induced by Cas9, Cas9^D^ or Cas9^H^ together with gTD1, gTD3 or gHR1b with scarless breakpoints and the proportions of this TD (bottom) in *Brca1^+/+^*and *Brca1^m/m^* TD reporter mouse ES cells. Scarless breakpoints are indicated by black arrows with indicated sgRNAs. The NGG PAMs are underlined. Letters with underline indicate PAM sequence. The underlined red “TGTC” is the MH sequence that mediates TD formation. The number of TDs with scarless breakpoints and the number of total TD events are shown in parenthesis after each proportion value in percentage.

In the formation of *RFP^+^* Inv II/Ter II TD products, however, Cas9 or Cas9n induces sequence alterations as “a scar” at the breakpoint in the sister chromatid template before allelic DNA synthesis, and the same scar could thus be introduced into the non-allelic *GFP* copy by allelic DNA synthesis, generating identical sequence alterations at the breakpoints in the tandem *GFP* copies of the TD products. Indeed, in each *RFP^+^* Inv II/Ter II TD product, the same indels were shared between the Inv II point and the Ter II point (Table EV 2 and Fig. EV5). Such *RFP^+^* Inv II/Ter II TD products comprised more than 50% of Cas9-induced TDs but only a small part or even none of Cas9n-induced TDs in both *Brca1^+/+^* cells and *Brca1^m/m^* cells (Fig. EV5). Surprisingly, “TGTC” was dominantly used as the MH junction of TDs in the TD reporter although the reason is unknown (Fig. EV5). Given the presence of two identical indel scar and one MH junction scar in RFP^+^ Inv II/ Ter II TD products, if these sites were not known as a Cas9 or Cas9n target, these three sites with a scar could be mistakenly identified as a breakpoint, leading to mischaracterization of the mechanisms underlying the TD formation.

Since many TD products induced by Cas9 and Cas9n in the TD reporter are *RFP^−^*, TD analysis could be biased by examining individual *RFP^+^* TD clones only. Thus, after TD was induced by Cas9 or Cas9n together with gTD1, gTD3 or gHR1b in TD reporter mouse ES cells, genomic DNA (gDNA) was isolated from a pool of cells containing both *RFP^+^* TD products and *RFP^−^* TD products. Using targeted PCR amplification with the primer pair Fa/Rb, we amplified the nested *GFP* sequences between *RFP* exon A and B from 100 ng gDNA for each treatment if TD products were generated as indicated (Fig. 3B). Two major bands were observed after PCR products were resolved by agarose gel electrophoresis. One with a larger size was designated as long TDs and the other with a smaller size as short TDs (Fig. 3C). We quantified the intensity of PCR bands and found that the intensity of PCR bands relative to the internal “*Gapdh*” PCR control from *Brca1^m/m^* cells was stronger than that from *Brca1^+/+^* cells, whether for TD induced by Cas9 together with gTD1, gTD3 and gHR1b or for TD induced by Cas9n together with gTD1, gTD3 and gHR1b (Fig. 3D). This is consistent with the role of *BRCA1* in suppression of TD formation.

To determine the breakpoints and junctions of TD products, the PCR bands corresponding to long TDs and short TDs were excised from the gel and subcloned for Sanger sequencing. Consistently, a small proportion of long TDs induced by Cas9 were the Inv I/Ter I type in both *Brca1^+/+^* cells and *Brca1^m/m^* cells and majority of Cas9n-induced long TDs were the same Inv I/Ter I products, all with a scarless breakpoint, the MH junction “TGTC” and an estimated ∼6.9-kb TD span (Fig. 3E). This indicates that the MH junction is not the breakpoint. The breakpoints in these TD products induced by Cas9 and Cas9n at different target sites in the TD reporter were masked due to lack of “scar” and would be hard to identify if they were not known as a target site for Cas9 or Cas9n.

After determining the proportions of long TDs with the Inv I/Ter I type in *Brca1^+/+^*cells and *Brca1^m/m^* cells (Fig. EV6A), we calculated the intensity of these long TDs induced by Cas9 or Cas9n by multiplying the combined intensity of TD PCR bands with the proportion of long TDs. The intensity of both Cas9-and Cas9n-induced long TDs relative to the internal PCR control “*Gapdh*” was enhanced by *Brca1* deficiency (Fig. EV6B). These results suggest that the frequency of long TDs with scarless breakpoints is stimulated in *Brca1^m/m^* cells as compared to *Brca1^+/+^* cells.

We also identified a single type of short TD products with 2,114-bp deletion between *RFP* exon A and B and with an estimate 4.78-kb TD span (Fig. EV6C). These TD products were detectable only for those induced by Cas9-gHR1b, Cas9^D^-gTD-1, Cas9^H^-gTD-1 and Cas9^H^-gHR1b in *Brca1^+/+^* cells and by Cas9-gTD-1, Cas9-gHR1b, Cas9^D^-gTD-3, Cas9^D^-gHR1b and Cas9^H^-gTD-3 in *Brca1^m/m^* cells (Fig. EV6C). It is likely that this type of TD products is generated as the Inv IV type of TD *via* the MMBIR mechanism (Fig. 6D).

### Brca1 suppresses nick-induced TD at natural genomic sites

As TD induced by a specific break in the TD reporter could be identified by PCR amplification of the invasion point or the termination point (Fig. 3B-D), we could thus similarly target any endogenous genomic sites for TD induction and TD analysis. We tested this possibility by inducing a site-specific break at the *H11*, *Rosa26* and *Col1a1* loci in mouse ES cells and amplifying the invasion point or the termination point of TD products using the outward primer pair F1/R1 (i.e., Primer Pair #1) or F2/R2 (i.e., Primer Pair #2) (Fig. 4A and EV7A). F1 and R1 or F2 and R2 are complementary to the sites upstream and downstream of the Cas9 or Cas9n cleavage sites, respectively; therefore, no PCR products should be produced with F1/R1 or F2/R2 without TD formation. Upon break induction by Cas9 or Cas9n, TD products are generated *via* aBIR-MMBIR or MMBIR, and PCR with F1/R1 or F2/R2 could amplify the TD region with either the invasion point represented by Inv I, Inv II, Inv III and Inv IV or the termination point represented by Ter I, Ter II, Ter III and Ter IV (Fig. 4A and EV7A). Other TD events might occur in a more complex form and are classified as complex events.

**Figure 4.**
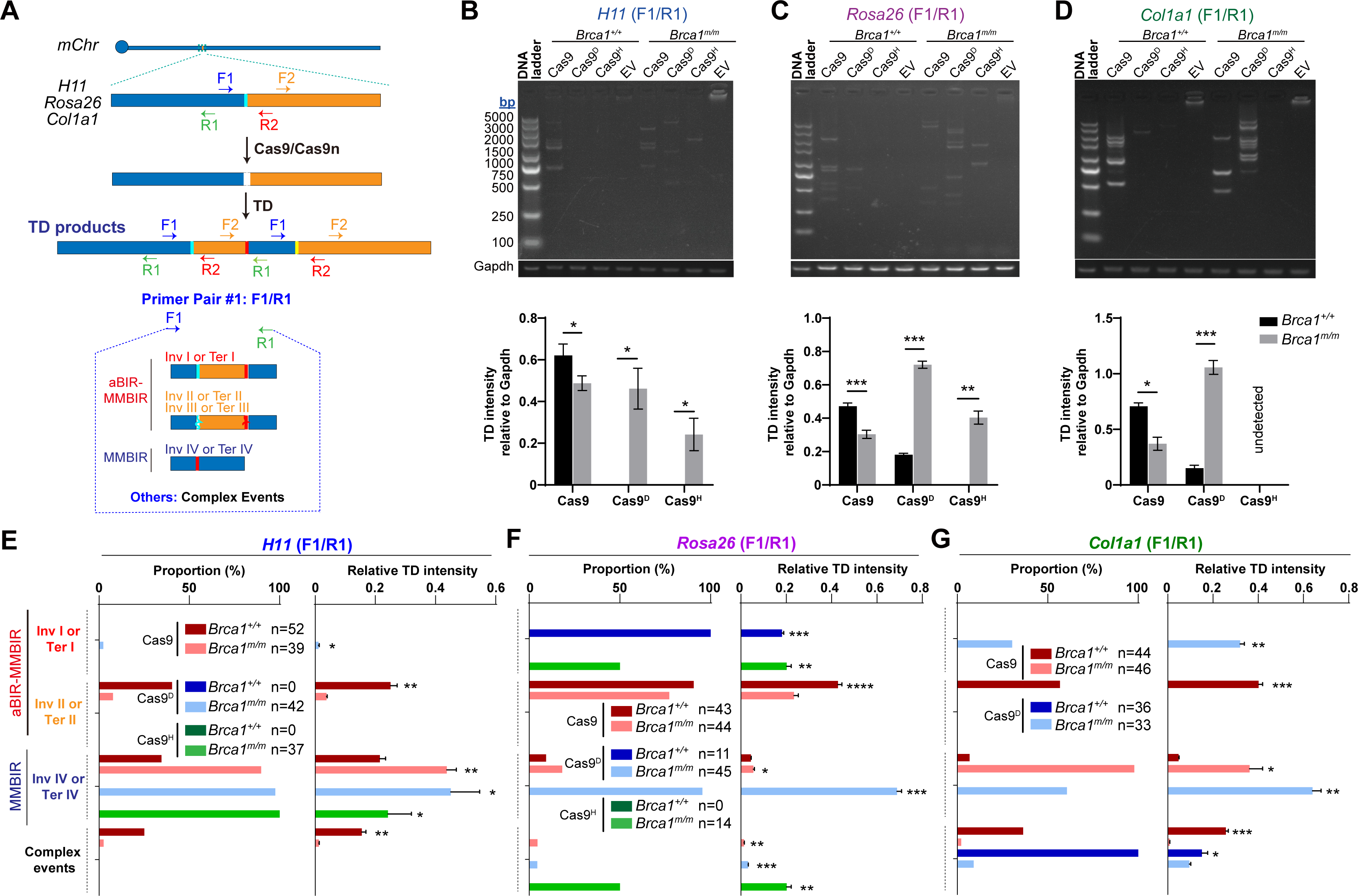
*Brca1* deficiency stimulates TDs induced by DNA nicks at natural sites. **(A)** Schematic for PCR-based detection of TDs induced by Cas9 and Cas9n at a natural site of the *H11*, *Rosa26* and *Col1a1* loci with F1/R1, an outward primer pair indicated in color arrows at the natural site. After site-specific TD induction by Cas9 or Cas9n, TD products shown can be amplified by PCR with F1/R1. TDs with Inv I or Ter I, Inv II or Ter II, Inv III or Ter III are considered as products of aBIR-MMBIR, and Inv IV or Ter IV as products of MMBIR. TDs in a more complex form are regarded as complex events. The green gap and the red gap denote the scarless breakpoints by Cas9 and Cas9n and the MH junction that mediates TD, respectively. The curve line in the green gap and the red gap indicates the presence of mutations. The outward primer pair F2/R2 is also indicated in color arrows at the original natural sites and the TD product. **(B-D)** Representative images (top) of PCR products with F1/R1 separated by DNA agarose gel electrophoresis for TDs induced by Cas9, Cas9^D^ or Cas9^H^ together with gH11 (**B**), gRosa26 (**C**) or gCol1a1 (**D**) and the intensity of these TDs in each lane relative to *Gapdh* (bottom) in *Brca1^+/+^* and *Brca1^m/m^*mouse ES cells. PCR products of a *Gapdh* region is used as internal PCR control. The TD intensity relative to *Gapdh* in the agarose gel is calculated as the ratio of the TD intensity to the *Gapdh* intensity measured by ImageJ software. **(E-G)** Distribution of TDs with each Inv/Ter type induced by Cas9, Cas9^D^ and Cas9^H^ at a natural site of the *H11* (**E**), *Rosa26* (**F**) and *Col1a1* loci (**G**) in *Brca1^+/+^* and *Brca1^m/m^* cells. The proportion of each Inv/Ter type in TDs (left) and the frequency of TDs with each Inv/Ter type (right) are shown. TDs with Inv I or Ter I and Inv II/Ter II are considered as products of aBIR-MMBIR, Inv IV or Ter IV as MMBIR, and others as complex events. Relative TD intensity for each Inv/Ter type is calculated as the TD intensity relative to *Gapdh* × the proportion of the Inv/Ter type in total TDs. The F1/R1 primer pair is also indicated for PCR detection of TDs. Column indicate the mean ± S.D. from three independent experiments in **B-D**, and statistical analysis is performed by paired two-tailed Student’s t test. In **E-G**, columns for relative TD intensity indicate the mean ± S.D. from three independent measurements of the TD intensity relative to *Gapdh*, and statistics is performed by paired two-tailed Student’s t test. n indicates the number of TD events analyzed in **E-G**. *, *P*<0.05; **, *P*<0.01; ***, *P*<0.001; ****, *P*<0.0001.

Using “*Gapdh*” PCR bands as the internal control, we found that PCR bands of Cas9-induced TDs with F1/R1 were less intense in *Brca1^m/m^*cells than in *Brca1^+/+^* cells, but those with F2/R2 were slightly stronger in *Brca1^m/m^* cells than in *Brca1^+/+^*cells (Fig. 4B-D and EV7B-D). Given that the same TD pools were amplified by F1/R1 and F2/R2, respectively, it is unexpected that the findings with F1/R1 are neither in line with those with F2/R2 nor consistent with the effect of *Brca1* deficiency on the formation of Cas9-induced TDs in the TD reporter (Fig. 2B, 4B-D and EV7B-D). This is possibly due to the difference in the annealing position of F1/R1 and F2/R2 in the genome to the break or the bias of primer pairs with different sequence contexts. Differently, PCR bands for Cas9n-induced TDs with F1/R1 or F2/R2 were weak or even absent in *Brca1^+/+^* cells but apparent in *Brca1^m/m^* cells (Fig. 4B-D and EV7B-D). The intensity of Cas9^D^ or Cas9^H^ -induced TDs relative to “*Gapdh*” was consistently strong in *Brca1^m/m^* cells but weak or even zero in *Brca1^+/+^* cells (Fig. 4B-D and EV7B-D), again indicating that *Brca1* suppresses nick-induced TD formation.

In order to further determine the effect of *Brca1* deficiency on each TD type induced by Cas9 or Cas9n at the *H11*, *Rosa26* and *Col1a1* loci of mouse ES cells, we subcloned mixed PCR products of genomic DNA containing site-specific TD with F1/R1 or F2/R2 to a TA clone vector and analyzed each subclone by Sanger sequencing. Cas9-induced TDs comprised of Inv II/Ter II, Inv IV/Ter IV and complex events whereas Cas9n-induced TDs included Inv I/Ter I, Inv II/Ter II, Inv IV/Ter IV and complex events (Fig. 4E-H and EV7E-H). As compared to *Brca1^+/+^* cells, *Brca1* deficiency consistently increased the level of Cas9-induced Inv IV/Ter IV TD (Fig. 4E-G and EV7E-G). In contrast, while Inv IV/Ter IV TDs induced by Cas9^D^ and Cas9^H^ at the three loci *H11*, *Rosa26* and *Col1a1* were not found in *Brca1^+/+^* cells, Inv IV/Ter IV TDs induced by Cas9^D^ at *H11*, *Rosa26* and *Col1a1* and by Cas9^H^ at *H11* were greatly elevated in *Brca1^m/m^*cells (Fig. 4E-G and EV7E-G). Similarly, Cas9^D^-induced Inv I/Ter I TDs at *H11* and *Col1a1* and Cas9^H^ -induced Inv I/Ter I TDs at *Rosa26* were undetectable in *Brca1^+/+^* cells but significantly stimulated by *Brca1* deficiency (Fig. 4E-G and EV7E-G). This analysis of each TD type again confirms that Brca1 suppresses nick-induced TD formation.

### The extent of involving aBIR in nick-induced TD varies between genomic sites

In the TD reporter, nick-induced *RFP^+^* TD formation stimulated by *Brca1* deficiency is triggered dominantly by allelic strand invasion but negligibly by MH-mediated non-allelic strand invasion in both *Brca1^+/+^* and *Brca1^m/m^* cells, suggesting the primary involvement of aBIR-MMBIR (Fig. 1D, 2D and 2E). As Cas9n-induced Inv IV/Ter IV and Inv I/Ter I TDs are mediated by MMBIR and aBIR-MMBIR, respectively, we wondered to what extent MMBIR and aBIR-MMBIR were differently involved in nick-induced TD formation at different sites in both *Brca1^+/+^* and *Brca1^m/m^*cells.

At the *H11* locus, MMBIR was the main responsible pathway for both Cas9-and Cas9n-induced TDs stimulated in *Brca1^m/m^* cells (Fig. EV8A and EV8B). At the *Rosa26* locus, both MMBIR and aBIR-MMBIR were used significantly for Cas9-induced TD formation (Fig. EV8C and EV8D). While stimulation of Cas9^D^-induced TDs in *Brca1^m/m^* cells were mostly mediated by MMBIR, Cas9^H^-induced TDs stimulated from zero in *Brca1^+/+^* cells to a significant level in *Brca1^m/m^*cells were either complex events or generated *via* aBIR-MMBIR (Fig. EV8C and EV8D). At the *Col1a1* locus, a significant level of Cas9-induced TDs in *Brca1^+/+^*cells were either complex events or generated by MMBIR and aBIR-MMBIR but MMBIR was responsible for most of Cas9-induced TDs in *Brca1^m/m^* cells. Cas9^D^-induced TDs at this site in *Brca1^+/+^* cells, despite at a low level, were also either complex events or generated by MMBIR (Fig. EV8E and EV8F); however, Cas9^D^-induced TDs, which were greatly elevated in *Brca1^m/m^* cells, were mostly generated *via* MMBIR (Fig. EV8E and EV8F). Cas9^H^ -induced TDs were absent in *Brca1^+/+^* cells but generated in *Brca1^m/m^* cells, and generation of these TDs were all mediated by MMBIR (Fig. EV8E and EV8F). These results indicate that the choice between MMBIR and aBIR-MMBIR in nick-induced TDs in both *Brca1^+/+^* and *Brca1^m/m^*cells vary significantly between genomic sites including the three endogenous loci (i.e., *H11*, *Rosa26* and *Col1a1*) and the TD reporter.

Of note, among complex TD events identified from the TD reporter and from the three natural sites *H11*, *Rosa26* and *Col1a1* by the primer pair F1/R1 and F2/R2, one Cas9^D^-induced complex TD event at *Rosa26* in *Brca1^m/m^* cells was generated by one-round non-allelic inverse strand invasion (Fig. EV9A). The remaining showed more than one round of template switching, up to 4 rounds, in both *Brca1^+/+^* cells and *Brca1^m/m^* cells (Fig. EV9B-D). We also found that three Cas9-induced TD events at *H11* in *Brca1^m/m^* cells were started by aBIR followed by MMBIR but terminated by end-joining with the second ends (Fig. EV9E).

### Template switching triggering MMBIR for nick-induced TD occurs mostly within 500 bp of allelic DNA synthesis

Upon encountering with DNA replication forks from either direction, Cas9n-induced DNA nicks are converted into one-ended DSBs with either a left end or a right end. Either of the left end and the right end can invade the allele of sister chromatid and is then extended by DNA synthesis. Subsequent displacement of nascent strand during allelic DNA synthesis could promote MH-mediated strand reinvasion into a non-allelic sequence of sister chromatid, generating identical TD products (Fig. 5A). Therefore, the TD formation *via* the aBIR-MMBIR pathway involves two rounds of DNA synthesis, first allelic for BIR and second non-allelic for template switching-directed MMBIR. We wondered how long the end was extended by allelic DNA synthesis prior to template switching (Fig. 5A). In a TD event generated by aBIR-MMBIR, allelic DNA synthesis of aBIR from the left end is undistinguishable from non-allelic DNA synthesis of MMBIR from the right end, and *vice versa* (Fig. 5A). Because the direction of the nick-colliding DNA replication fork is unknown, allelic DNA synthesis could not be assigned so that the length of allelic DNA synthesis could not be determined. However, it was well documented that nascent DNA strands are frequently displaced for template switching in initial homology-mediated BIR whereas later DNA synthesis is much more extended even in MMBIR (Liu & Malkova, 2022; Sakofsky *et al*, 2015; Smith *et al*, 2007; Anand *et al*, 2014). We thus considered shorter length of DNA synthesis as allelic and the other as non-allelic in a given TD product generated by aBIR-MMBIR.

**Figure 5.**
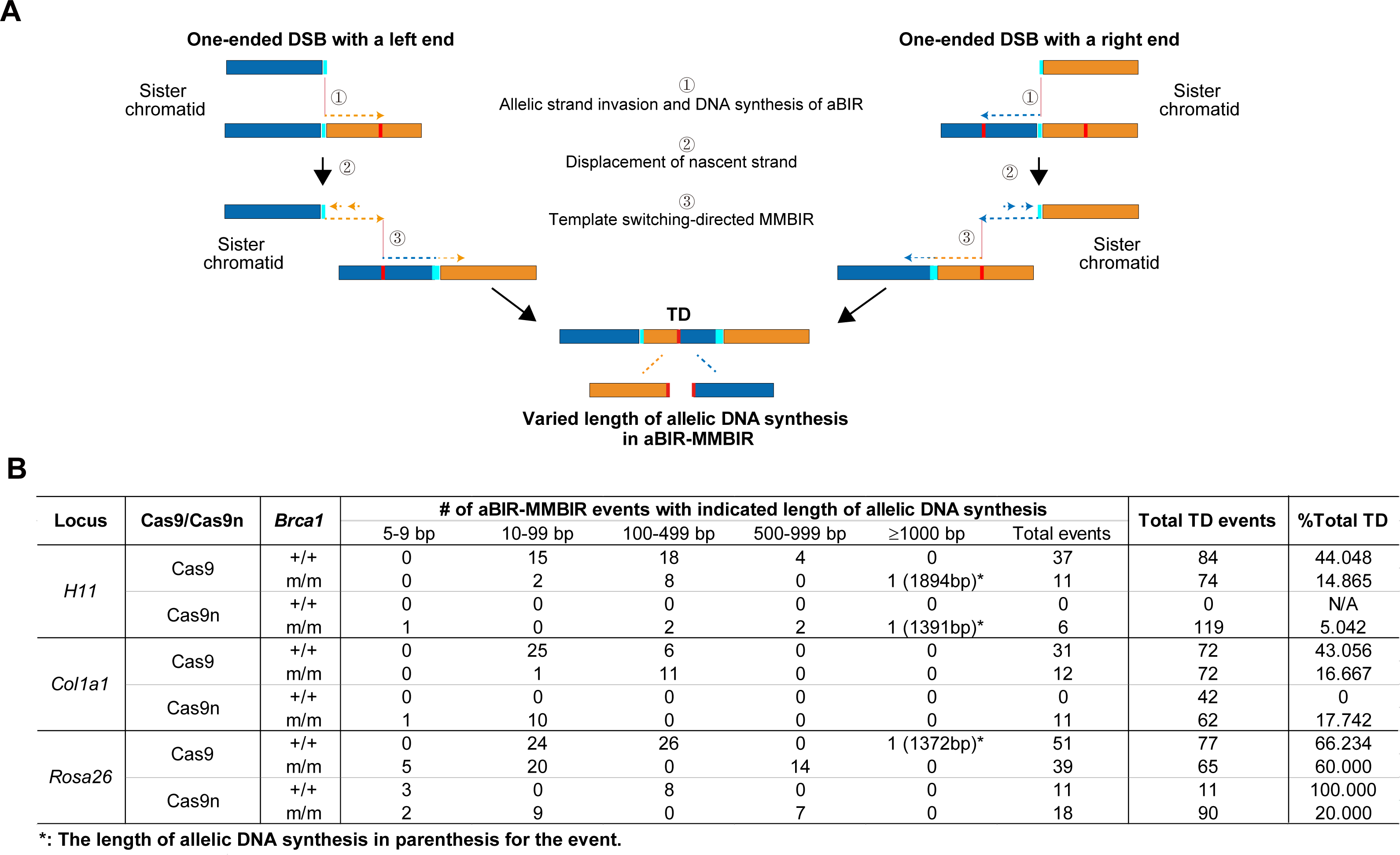
Break-induced allelic DNA synthesis extends in limited distance from the breakpoints prior to template switching. **(A)** Schematic for aBIR-MMBIR-mediated TD formation induced by one-ended DSBs with a left end or a right end. Upon one-ended DSBs, the left end or the right end of the DSBs invades the allele of sister chromatid and extends DNA synthesis. However, allelic DNA synthesis could be disrupted at a random distance from the invasion point, generating varied length of nascent DNA. Displaced DNA strand could thus revert and reinvade non-allelic region of sister chromatid with MH, triggering template switching-directed MMBIR. TD products generated from the left end could be identical to those from the right end in this model. **(B)** The length of allelic DNA synthesis prior to template switching in aBIR-MMBIR TD events induced by Cas9 and Cas9n at the natural *H11*, *Col1a1* and *Rosa26* site.

Of 37 aBIR-MMBIR TD events induced by Cas9 at *H11*, 31 at *Col1a1* and 51 at *Rosa26* in *Brca1^+/+^*cells, 33, 31 and 50 had allelic DNA synthesis less than 500 bp, respectively (Fig. 5B). Allelic DNA synthesis also extended less than 500 bp in most of aBIR-MMBIR TD events induced by Cas9 in *Brca1^m/m^* cells (Fig. 5B). While aBIR-MMBIR TD events induced by Cas9n were absent at *H11* and *Col1a1* in *Brca1^+/+^* cells, a half of 6 aBIR-MMBIR TD events induced by Cas9n at *H11* and all of 11 at *Col1a1* had allelic DNA synthesis less than 500 bp in *Brca1^m/m^* cells (Fig. 5B). Similarly, allelic DNA synthesis at the *Rosa26* locus prior to template switching was less than 500 bp in all of 11 aBIR-MMBIR TD events induced by Cas9n in *Brca1^+/+^* cells and 11 out of 18 in *Brca1^m/m^* cells (Fig. 5B). These results support that template switching for TD formation induced by two-ended DSBs or replication-coupled one-ended DSBs occur within a limited distance upstream of the site of allelic strand invasion in mammalian cells.

### Short TDs induced at a given site by Cas9 and Cas9n are heterogenous

At a given site where a TD product is formed, the position of MH-mediated strand invasion and the extent of DNA synthesis may differ, generating TDs with various span size and different MH usage. As a significant level of TDs can be induced by Cas9 in both *Brca1^+/+^* cells and *Brca1^m/m^* cells, we also wondered whether *Brca1* played a role in controlling the span size of these TDs. In the TD reporter, nearly 100% of *RFP^+^* TDs induced by Cas9 were Inv II/Ter II in both *Brca1^+/+^* cells and *Brca1^m/m^* cells. These TDs generated *via* aBIR-MMBIR were estimated to span ∼6.9 kb (Fig. 1A and 2C-E), giving no indication on such a role of *Brca1*. While TD span sizes were more heterogenous at the *H11*, *Rosa26* and *Col1a1* loci for Cas9-induced TDs in either *Brca1^+/+^* cells or *Brca1^m/m^* cells, e.g., 0.34-5.39 kb at *H11*, 0.59-5.19 kb at *Rosa26* and 0.29-5.76 kb at *Col1a1* (Fig. EV10A and EV10B), we found no consistent effect of *Brca1* deficiency on the span sizes of these TDs.

Span sizes of Cas9n-induced TDs were also heterogenous at the *H11*, *Rosa26* and *Col1a1* loci in either *Brca1^+/+^* cells or *Brca1^m/m^* cells, e.g., 0.27-3.90 kb at *H11*, 0.56-9.70 kb at *Rosa26* and 0.25-7.28 kb at *Col1a1* for Cas9^D^-induced TDs, and 0.88-2.95 kb at *H11*, 0.74-2.90 kb at *Rosa26* and 0.40-0.74 kb at *Col1a1* for Cas9^H^-induced TDs (Fig. EV10A and EV10B). When Cas9^D^-induced TDs were detectable at *Rosa26* by PCR with F1/R1 and at *Col1a1* by PCR with F2/R2 in *Brca1^+/+^*cells, the span sizes of these TDs were increased by *Brca1* deficiency (Fig. EV10A and EV10B). However, due to general lack of Cas9n-induced TDs in *Brca1^+/+^* cells, it was difficult to confirm a role of *Brca1* in controlling the span size of Cas9n-induced TDs.

The use of MH was expected in both MMBIR and aBIR-MMBIR that generate *BRCA1*-linked TDs. We wondered whether the type of MH involved varied between these two pathways. In the TD reporter, “TGTC” and “CAGG” were the only MH found in TDs generated by aBIR-MMBIR and by MMBIR, respectively (Fig. 3A, 3E, EV5 and EV6C). In contrast, the MH type was more heterogeneous at the junction of TDs induced by Cas9 and Cas9n at the *H11, Rosa26* and *Col1a1* sites in both *Brca1^+/+^* cells and *Brca1^m/m^* cells whether TDs were generated *via* MMBIR or aBIR-MMBIR. In TDs generated by aBIR-MMBIR, the length of MH varied between 0 bp to 12 bp for Cas9-induced TDs and between 0 bp to 11 bp for Cas9n-induced TDs at these three sites (Fig. EV11A-C). In contrast, in those TDs generated by MMBIR, the length of MH varied between 0 bp to 8 bp for both Cas9 and Cas9n-induced TDs at these three sites (Fig. EV11D-F). While it was surprising that MH was not found in several TD events (Fig. EV11A-F), this is in line with previous observation that non-allelic strand invasion including template switching can occur, albeit infrequently, without assistance of homology or MH (Sakofsky *et al*, 2015).

### RAD51 loading by BRCA1 is dispensable for BRCA1-mediated TD suppression

Previous study has shown that disruption of BRCA1 BRCT domains destabilized BRCA1 proteins, simultaneously inactivating both functions of BRCA1 in end resection and RAD51 loading in HR(Williams *et al*, 2003). In BRCA-mediated TD suppression, it remains unknown which function of *BRCA1* is exactly required for TD suppression. We thus depleted *Rad51* or *Brca2* in TD reporter mouse ES cells and analyzed the effect of Rad51 loading on TD suppression. As both *Rad51* and *Brca2* were efficiently depleted by siRNAs (Fig. EV12A and EV12B), TDs induced by Cas9 and Cas9n at all six sites of the TD reporter were not elevated by depletion of *Rad51* or *Brca2*; instead, depletion of *Rad51* or *Brca2* reduces Cas9n-induced TDs (Fig. 6A and EV12C-E).

**Figure 6.**
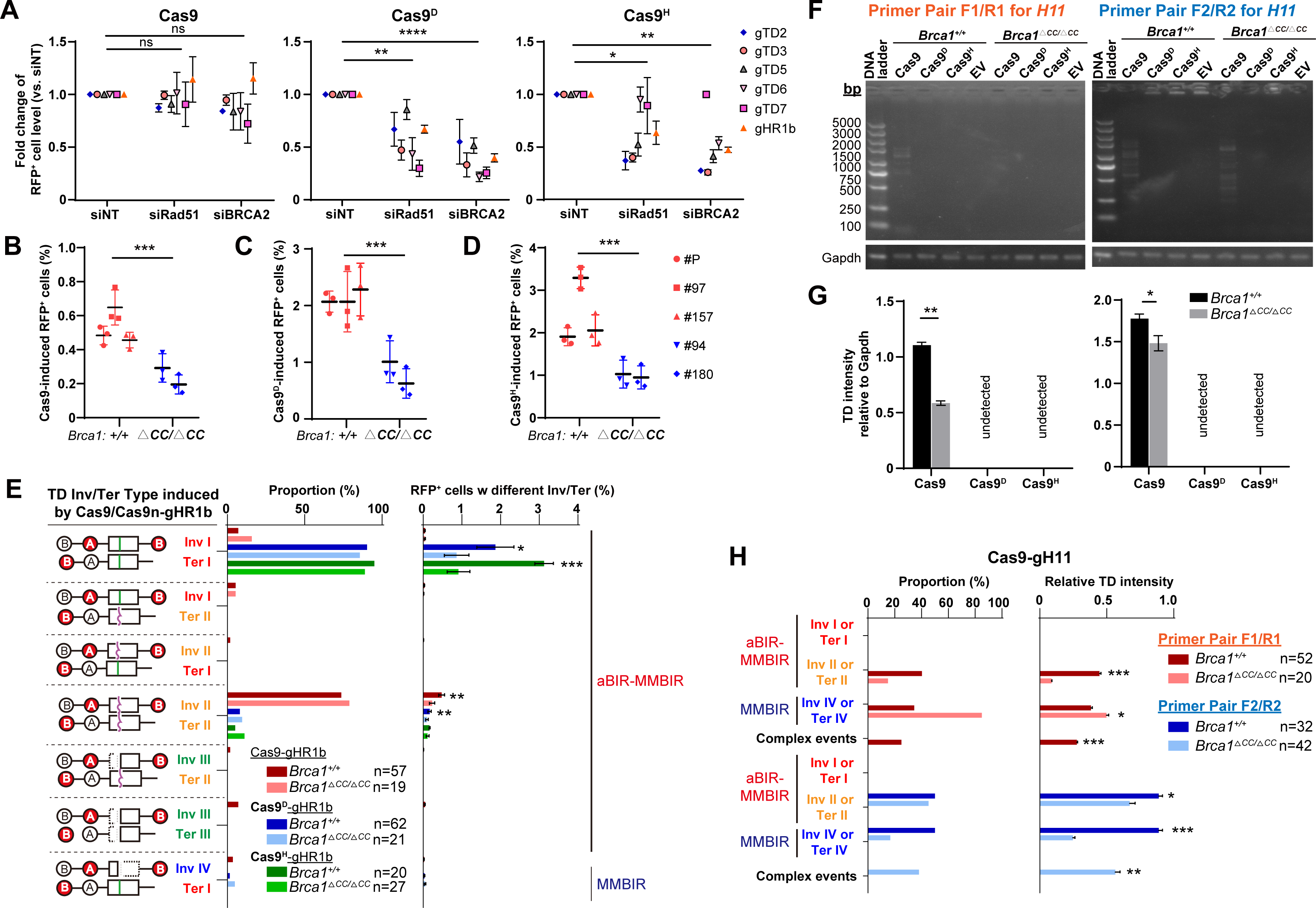
Rad51 loading is dispensable for Brca1-mediated TD suppression. **(A)** Effect of *Rad51* depletion and *Brca2* depletion on induction of *RFP^+^* cells by Cas9 (left), Cas9^D^ (middle) or Cas9^H^ (right) together with 7 sgRNAs indicated by different shapes. Each color shape with an error bar represents the mean ± S.D. of three independent experiments for one sgRNA. Fold change of *RFP^+^* cell frequencies with siRad51 or siBrca2 treatment is calculated by normalizing the percentages of *RFP^+^* cells induced by Cas9, Cas9^D^ and Cas9^H^ with siNT treatment to 1. siNT: non-target siRNA as negative control; siRad51 and siBrca2: siRNA against *Rad51* and *Brca2*, respectively. **(B-D)** Frequencies of *RFP^+^* cells induced by Cas9 (**B**), Cas9^D^ (**C**) or Cas9^H^ (**D**) together with gHR1b in three *Brca1^+/+^* clones and two *Brca1^ΔCC/ΔCC^* clones. Each color shape with an error bar represents the mean ± S.D. of three independent experiments with one clone. **(E)** Distribution of *RFP^+^* cells with each Inv/Ter type induced by Cas9, Cas9^D^ and Cas9^H^ together with gHR1b in *Brca1^+/+^*and *Brca1^ΔCC/ΔCC^* cells. The proportion of each Inv/Ter type in *RFP^+^* cells and the frequency of *RFP^+^*cells with each Inv/Ter type are shown. *RFP^+^* cells with Inv I/Ter I, Inv I/Ter II, Inv II/Ter I, Inv II/Ter II, Inv III/Ter II and Inv III/Ter III are classified as products of aBIR-MMBIR, and Inv IV/Ter I and Inv IV/Ter II as MMBIR. The frequency of *RFP^+^* cells with each Inv/Ter type is calculated as the overall frequency of *RFP^+^* cells × the proportion of the Inv/Ter type in *RFP^+^*cells. **(F and G)** Representative images (**F**) of PCR products separated by DNA agarose gel electrophoresis for TDs induced by Cas9, Cas9^D^ or Cas9^H^ at a natural site of *H11* and the intensity of these TDs in each lane shown relative to *Gapdh* (**H**) in *Brca1^+/+^* and *Brca1^ΔCC/ΔCC^* mouse ES cells. The primer pair F1/R1 (left) and F2/R2 (right) are indicated for PCR products of TDs (**F**). PCR products of a *Gapdh* region is used as internal PCR control. The TD intensity relative to *Gapdh* in the agarose gel is calculated as the ratio of the TD intensity to the *Gapdh* intensity measured by ImageJ software. **H,** Distribution of TDs with each Inv/Ter type induced by Cas9-gH11 at *H11* in *Brca1^+/+^* and *Brca1^ΔCC/ΔCC^* cells. TDs were identified by PCR with the primer pair F1/R1 and F2/R2 as indicated. The proportion of each Inv/Ter type in TDs (left) and the frequency of TDs with each Inv/Ter type (right) are also shown. TDs with Inv I or Ter I and Inv II/Ter II are considered as products of aBIR-MMBIR, Inv IV or Ter IV as MMBIR, and others as complex events. Relative TD intensity for each Inv/Ter type is calculated as the TD intensity relative to *Gapdh* × the proportion of the Inv/Ter type in total TDs. Each symbol with an error bar represents the mean ± S.D. of three independent experiments for one sgRNA in **A** and for one isogenic clone in **B-D**. Statistics is performed by two-tailed Student’s t testing in **A** and by the Mann–Whitney test in **B-D**. Columns indicate the mean ± S.D. from three independent measurements in **E, G** and **H**, and n indicates the number of TD events analyzed in **G**. Statistical analysis is performed by paired two-tailed Student’s t test in **E, G** and **H**. *, *P*<0.05; **, *P*<0.01; ***, *P*<0.001.

It is well-established that RAD51 loading by BRCA1 is mediated by the coiled-coil (CC) domain of BRCA1 (Sy *et al*, 2009). Thus, to further determine the role of Rad51 loading in Brca1-mediated TD suppression, we deleted part of exon 12 encoding the CC domain of Brca1 in TD reporter mouse ES cells by paired Cas9 approach (Fig. EV12A) (Guo *et al*, 2018), disabling the activity of the CC domain and generating *Brca1^△CC/△CC^* TD reporter mouse ES cells defective in Rad51 loading. 2 *Brca1^△CC/△CC^* TD reporter mouse ES clones as well as 3 isogenic *Brca1^+/+^* TD reporter mouse ES clones were established and validated by Sanger sequencing (Fig. EV13A). The protein level of Brca1 with the truncated CC domain was comparable to that of wild-type Brca1 (Fig. EV13B); however, Rad51 loading upon treatment with the PARP inhibitor Olaparib was indeed defective in *Brca1^△CC/△CC^* TD reporter mouse ES clones (Fig. EV13C), consistent with previous finding (Nacson *et al*, 2020). TDs induced by Cas9 and Cas9n in the TD reporter were not elevated, but reduced in 2 isogenic *Brca1^△CC/△CC^* clones as compared to 3 isogenic *Brca1^+/+^*clones (Fig. 6B-D), indicating that Rad51 loading is not required for *Brca1*-mediated TD suppression. In contrast, Rad51 loading by Brca1 might promote TD formation although the underlying mechanism is yet to be identified.

To determine whether the TD types are altered between *Brca1^+/+^* cells and *Brca1^△CC/△CC^* cells, *RFP^+^* TD cells were sorted and individual *RFP^+^* TD clones were analyzed for the sequences of the invasion point and the termination point in TD products by PCR amplification followed by Sanger sequencing. Majority of Cas9-and Cas9n-induced *RFP^+^* TDs respectively remained Inv II/Ter II and Inv I/Ter I, both of which were reduced in *Brca1^△CC/△CC^* cells (Fig. 6E). Without sorting for *RFP^+^* cells, a pool of *RFP^+^* and *RFP^−^* cells were also used to determine TD formation by PCR after TD induction by Cas9 and Cas9n in the TD reporter (Fig. EV13D). The intensity of TDs relative to the internal “*Gapdh*” PCR control was weaker in *Brca1^△CC/△CC^* cells than in *Brca1^+/+^* cells (Fig. EV13E), again excluding any requirement of Rad51 loading for Brca1-mediated TD suppression.

Further, we analyzed TD formation induced by Cas9 and Cas9n at the endogenous *H11* site in both *Brca1^+/+^* cells and *Brca1^△CC/△CC^* cells and found that the intensity of PCR bands for Cas9-induced TDs relative to the internal “*Gapdh*” PCR control was stronger in *Brca1^+/+^* cells than in *Brca1^△CC/△CC^* cells (Fig. 6F and 6G). Cas9-induced TDs contained Inv II/Ter II, Inv IV/Ter IV and complex TD events, and the Inv II/Ter II TD type was reduced in *Brca1^△CC/△CC^* cells (Fig. 6H, EV13F and EV13G). More importantly, while *Brca1* deficiency was expected to stimulate Cas9n-induced TDs at the endogenous sites, no PCR bands were found for Cas9n-induced TDs either in *Brca1^+/+^*cells or in *Brca1^△CC/△CC^* cells (Fig. 6F and 6G), further confirming that Rad51 loading through the CC domain of Brca1 is dispensable for Brca1-mediated TD suppression.

## Discussion

Human *BRCA1*-deficient tumors accumulate MH-mediated TDs ranging from 1.6 kb to 51 kb (median value of 11 kb) (Menghi *et al*, 2018). Several mechanisms have been proposed to explain the formation of TDs associated with *BRCA1* deficiency (Willis *et al*, 2017; Feng *et al*, 2022; Kamp *et al*, 2020). It is expected that these underlying mechanisms may differ in response to different types of endogenous DNA lesions in *BRCA1*-deficient cells. Using a HR reporter containing two tandem GFP repeats in *BRCA1*-deficient cells, our previous study has suggested an increased involvement of aborted allelic HR followed by template switching-directed homology-mediated BIR in TD formation induced by DNA nicks, not by classical two-ended DSBs (Feng *et al*, 2022). However, this mechanism may not be applicable to over 50% of the genome that lacks neighboring repeats. Here, we analyzed the formation of nick-induced TDs in a repeat-less TD reporter or in a natural genomic region containing no neighboring homology and found that, in addition to the MMBIR pathway alone, aBIR terminated prematurely by template switching-directed MMBIR (i.e., aBIR-MMBIR) also served as a major mechanism to promote nick-induced TDs in *Brca1*-deficient cells (Fig. EV14). In addition, in line with Cas9n-induced on-target rearrangements previously reported (Feng *et al*, 2022), a significant level of on-target TDs induced by Cas9n could even exceed 3% in normal (*BRCA1*-proficient) cells and 4% in *BRCA1*-deficient cells, again demonstrating a source of safety concerns and a need to resolve such concerns in Cas9n-based applications in base editing and prime editing, two of major genome editing technologies (Chen & Liu, 2023). This is at least in part due to long target residency of Cas9n after DNA nicking (Feng *et al*, 2022, 2021; Liu *et al*, 2022). This type of on-target chromosomal rearrangements should be taken into consideration in safety assessment of genome editing for clinical application.

Upon colliding with DNA replication forks, DNA nicks are converted into one-ended DSBs. It is well-established in yeast that one-ended DSBs primarily engage BIR for repair whereas the most common HR pathway SDSA is preferred over BIR in repair of classical two-ended DSBs unless homology at either ends of the DSBs are short or Mph1 is deficient (Liu & Malkova, 2022; Anand *et al*, 2013; Llorente *et al*, 2008; Wu & Malkova, 2021; Malkova *et al*, 2005; Mehta *et al*, 2017; Prakash *et al*, 2009; Stafa *et al*, 2014; Piazza *et al*, 2019; Pham *et al*, 2021). In mammalian cells, studies have also supported preferential use of BIR in repair of one-ended DSBs arising from replication fork stalling, collision of DNA nicks with replication forks, or erosion of telomeres (Feng *et al*, 2022; Willis *et al*, 2014; Costantino *et al*, 2014; Li *et al*, 2021; Dilley *et al*, 2016; Bhowmick *et al*, 2016; Hu *et al*, 2019; Minocherhomji *et al*, 2015; Zhang *et al*, 2023; Lydeard *et al*, 2007). After end resection, one-ended DSBs derived from nicks most likely invade the allele of the sister chromatid and trigger allelic BIR for optimal repair fidelity (Liu & Malkova, 2022; Anand *et al*, 2013; Llorente *et al*, 2008; Wu & Malkova, 2021). The other ends of the nicks are left on the sister chromatid and could be sealed by DNA ligases, giving an intact sister chromatid template for allelic BIR (Fig. EV14). However, although it is likely infrequent, the converging forks from the other direction could encounter the nicks before the other ends of the nicks are ligated, generating additional dsDNA ends to form two-ended DSBs. As these two-ended DSBs are coupled with DNA replication, it is possible that allelic BIR remains the major pathway over allelic SDSA in HR repair of these DSBs. However, unlike replication-coupled nick-induced TD that is consistently suppressed by BRCA1, TD induced by classical two-ended DSBs was not always stimulated by *BRCA1* deficiency, but even reduced, possibly owing to the sequences and chromatin context of the TD site or other unknown reasons. In addition, *Brca1* deficiency did not stimulate TDs induced by conventional two-ended DSBs in the SCR-RFP reporter and the Tus-Ter SCR-RFP reporter (Willis *et al*, 2017; Feng *et al*, 2022); however, *GFP* repeats in these reporters could interfere Cas9-induced TD formation represented by *GFP^−^RFP^+^*cells. Therefore, we could not yet draw a conclusion on the contribution of classical two-ended DSBs to the enrichment of *BRCA1*-linked TD in a genome region containing no neighboring homology.

In either BIR or SDSA, one-ended DSBs would normally undergo 5′-to-3′ end resection, creating a single 3’ ssDNA end for the assembly of a RAD51 filament (Liu & Malkova, 2022; Cejka & Symington, 2021). The filament could thus search homologous sequences and invade a homologous template for D-loop displacement synthesis, extensive in BIR or short-patched in SDSA (Liu & Malkova, 2022; Cejka & Symington, 2021). While the homologous template is primarily provided by the homologous allele in sister chromatid, allowing conservative DNA synthesis for allelic BIR or allelic SDSA, non-allelic homology-mediated strand invasion could lead to aberrant chromosomal rearrangements. However, 5’-to-3’ end resection is inefficient in *BRCA1*-deficient cells and the length of the RAD51 filament that is subsequently formed may not be sufficient, prohibiting productive search for homology or invasion into a homologous template. Thus, this end could increasingly engage MH-mediated strand invasion into a site on either side of the original nick at the sister chromatid template and promote MMBIR, generating either TDs or deletions associated with *BRCA1* deficiency.

In addition, in this current study, we detected a significant level of allelic DNA synthesis followed by template switching in TD products associated with *Brca1* deficiency. This suggests that allelic strand invasion with homology pairing remains robust in *BRCA1*-deficient cells but nascent strand displacement and template switching occur more frequently after allelic DNA synthesis. In the absence of second ends, displaced nascent strand could then either re-invade allelic homologous template or engage MH-mediated re-invasion into a site on either side of the original nicking site at the sister chromatid template. The template switching in the latter case leads to generation of MMBIR-mediated TDs or deletions associated with *BRCA1* deficiency. Strand annealing or end-joining with the second ends could complete repair if second ends are available, preventing extensive DNA synthesis. However, due to inefficient end resection of the second ends in *BRCA1*-deficient cells, strand annealing with second ends is defective. Consequently, this could increase the probability of template switching for displaced nascent strand to induce MMBIR-mediated TDs (Fig. EV14).

In budding yeast, template switching occurs at frequencies as high as 20% of BIR events and mostly within the first 5 kb of the strand invasion site (Smith *et al*, 2007; Anand *et al*, 2014). Template switching also arises at a rate of ∼10^-6^ from SDSA-mediated mate-type switching of budding yeast (Hicks *et al*, 2010), but this rate is nearly 10^5^-fold lower than that of template switching in BIR. In the current study, the frequencies of TD containing the signature of template switching are about 0.2-1.5% for two-ended DSBs induced by Cas9 and about 1-3% for one-ended DSBs induced by Cas9 nickases at the TD reporter lacking neighboring repeats as a homologous template. The high frequencies of TDs are consistent with the notion that TDs are not a mutational outcome of SDSA, but a product of MMBIR or aborted aBIR. Previous studies have demonstrated that template switching occurs more frequently in close proximity to the site of the prior strand invasion (Smith *et al*, 2007; Anand *et al*, 2014). It is also expected that template switching between proximal regions of sister chromatids is the most preferable. This could explain the high frequencies of TDs generated by MH-mediated re-invasion of displaced nascent strand from allelic DNA synthesis into a sequence near the original nicking site of the sister chromatid template. As the largest TD span size detected here is 9.7 bp, this indicates that displaced nascent strands re-invade a region less than 5 kb from allelic strand invasion site at the sister chromatid template. Also, in TD products found in *Brca1^+/+^*and *Brca1^m/m^* mouse ES cells, the length of displaced nascent strands from allelic DNA synthesis varies between 5 bp and 1,894 bp, but mostly within 500 bp, prior to template switching, suggesting possible involvement of non-processive DNA polymerases in allelic DNA synthesis that may be prone to nascent strand displacement in the extending D-loop (Liu & Malkova, 2022; Smith *et al*, 2007).

As shown here, aBIR-MMBIR involving template switching is one of the major pathways for stimulation of nick-induced TD in *Brca1*-deficient cells. How does BRCA1 suppress template switching in BIR? Previous work has indicated that the displaced invading strand tends to re-invade if the other end of the break is absent or defective for strand annealing that also requires end resection (Scully *et al*, 2019; Chandramouly *et al*, 2013; Liu & Malkova, 2022; Smith *et al*, 2007). In the presence of second ends, inefficient resection of second ends due to *BRCA1* deficiency could delay strand annealing and therefore promote template switching of the displaced invading strand during allelic BIR or allelic SDSA, increasing the TD frequency. Indeed, in repair of two-ended DSBs, defective resection of a second end in *BRCA1*-deficient cells delay the capture of the second end for gene conversion (GC) termination by SDSA, drive re-invasion of the displaced strand and further extend DNA synthesis, shifting the bias of GC towards LTGC, a BIR-like process (Feng *et al*, 2022; Chandramouly *et al*, 2013; Willis *et al*, 2014). On the other hand, collision of nicks with converging forks from one direction is expected to occur much more frequently than that from both directions, which is required to convert the nicks into two-ended DSBs. If generation of one-ended DSBs is more likely, it is possible that *BRCA1* deficiency could slow converging forks due to the role of *BRCA1* in restart of stalled forks in response to replication stress (Taglialatela *et al*, 2017; Schlacher *et al*, 2012; Deshpande *et al*, 2022). Consequently, this could reduce collision of the nicks with converging forks from both directions or delay the merging of extended D-loop with a converging fork in repair of one-ended DSBs by allelic BIR (Fig. EV14), thus increasing the chances of template switching for the displaced invading strand. As FANCM is found to catalyze displacement of the nascent strand from D-loop (Piazza *et al*, 2019; Sun *et al*, 2008), BRCA1 may play a role in suppressing this process, preventing template switching that is simulated by displaced invading strand. However, this possibility has yet to be tested.

Not every key function of *Brca1* is required for TD suppression. Deletion of the CC domain does not stimulate enrichment of TD products associated with *Brca1* deficiency. Similarly, TD was not stimulated, but modestly reduced, by depletion of *Rad51* or *Brca2*, consistent with little accumulation of this structural variation (SV) type in *BRCA2*-deficient tumors (Menghi *et al*, 2018). These results suggest that unlike the other BRCA1 functions, RAD51 loading by BRCA1 may not help suppress TD formation induced by DNA nicks, but rather mediate. In contrast, it is believed that BRCA1-mediated end resection played a major role in suppressing TD formation. As both BRCA1 functions are required for the formation of the RAD51 filaments for homology search in BIR, it is paradoxical that the TD formation is oppositely regulated by RAD51 loading and end resection. However, because RAD51 loading and end resection are two neighboring steps leading to the presynaptic assembly of the RAD51 filaments, it is apparent that the defect in each step causes a different outcome in RAD51 filament formation. BRCA1-mediated end resection generates 3’-ssDNA of sufficient length, allowing the use of a sufficiently long RAD51 filament for homology search in BIR. Differently, RAD51 loading ensures the sufficient RAD51 density in the fully assembled RAD51 filament at 3’-ssDNA of given length, short or long dependently of end resection, for HR including BIR. Therefore, the length of the RAD51 filament may determine the choice between allelic BIR and MMBIR or the choice between the D-loop extension of allelic BIR and the D-loop disruption followed by template switching.

As previously suggested, template switching may be facilitated by the use of DNA polymerases that are less processive (Liu & Malkova, 2022; Sakofsky *et al*, 2015; Northam *et al*, 2014). It remains unclear whether the length of homology pairing following allelic strand invasion determine the engagement of processive or less processive DNA polymerases for D-loop DNA synthesis. If so, due to inefficient end resection that generates only a short length of homologous 3’-ssDNA in *BRCA1*-deficient cells, a short RAD51 filament formed could engage a less processive DNA polymerase such as translesion synthesis (TLS) polymerase Polz, leading to more disruption of D-loop extension and more frequent displacement of nascent strand. Studies in *Drosophila* and yeast have shown that replicative and TLS polymerases could compete for initiation of DNA synthesis during HR that includes SDSA and BIR (Sakofsky *et al*, 2015; Kane *et al*, 2012; Deem *et al*, 2011). In particular, interruption of homology-driven Polδ-catalyzed BIR could trigger template switching involving 0-6 nt of MH, inducing MMBIR directed by TLS polymerases Polζ and Rev1 (Sakofsky *et al*, 2015). Therefore, the end that is not sufficiently resected may invade the allelic site in the sister chromatid, but unlike a fully resected end, preferably engages a DNA polymerase such as TLS polymerases that might be more susceptible to dissociation from DNA synthesis (Sakofsky *et al*, 2015; Kane *et al*, 2012; Deem *et al*, 2011). As a result, during allelic BIR, nascent DNA strand could be prematurely displaced from extended D-loop, increasing the opportunity of template switching into a site at the sister chromatid for MMBIR-mediated TD. In contrast, it is possible that that the density of RAD51 in the RAD51-3’-ssDNA filament, regardless of the 3’-ssDNA length, has no such effect on selecting DNA polymerases, but is only limited to homology search and initiation of DNA synthesis. While it is reasonable to predict that *BRCA1*-linked TDs would not accumulate in tumors carrying pathogenic mutations of the CC domain, these possible mechanisms warrant further investigation.

In cancer genome analysis, knowing the exact point of a DNA break in the genome is not only critical for revealing the process of the DNA breakage and the type of original DNA lesion, but could also help elucidate the mechanisms by which break-induced chromosomal rearrangements are initiated, extended and terminated (Alkan *et al*, 2011; Li *et al*, 2020; Conrad *et al*, 2010; Lam *et al*, 2010; Malhotra *et al*, 2013; Cheloshkina & Poptsova, 2021). Inaccurate identification of a breakpoint would potentially lead to mischaracterization of DNA lesions and erroneous deduction of the underlying mechanisms for cancer-specific structural variations. Currently, a breakpoint is often defined as a junction characterized with the presence of sequence alteration for break-induced structural variations (Conrad *et al*, 2010; Lam *et al*, 2010; Malhotra *et al*, 2013; Quinlan *et al*, 2010). However, as revealed in this study, allelic DNA synthesis following allelic strand invasion restores the sequence of the breakpoint by Cas9 or Cas9n and shifts the MH-mediated junctions away from the breakpoint in break-induced TD. If these breakpoints were not known in advance as Cas9 or Cas9n target sites, the MH-mediated junctions could be conveniently mis-regarded as the breakpoints and allelic strand invasion followed by template switching could be missed as one of the major mechanisms for *BRCA1*-linked TD. It is therefore possible that equaling a junction to a breakpoint in the TD products could result in incorrect identification of the breakpoints in cancer genomes and misunderstanding of the mechanisms for break-induced chromosomal rearrangements.

Prior to this study, the use of DNA repeat-based reporters helped reveal the mechanisms by which *BRCA1*-linked TD is formed in response to stalled replication forks or DNA nicks (Willis *et al*, 2017; Feng *et al*, 2022). These TD mechanisms may be applicable to a genome region containing neighboring repeats but do not represent those in the genome lacking neighboring homology. Given that ∼50% of the human genome is repeat-less (Lander *et al*, 2001; Venter *et al*, 2001), it is important to determine the underlying mechanisms for the TD formation in these repeat-less region of the genome. To this end, this study generated a repeat-less TD reporter directly from the SCR-RFP reporter integrated in the genome of mouse ES cells, developed a TD assay at endogenous sites lacking tandem homologous repeats to analyze TD products induced by DNA nicks, the most frequent type of endogenous DNA lesions(Caldecott, 2022), and revealed different frequency and regulation of TD formation between genomic regions with or without neighboring repeats. Therefore, the findings of this study not only further advance our understanding of the TD formation in cancer genomes associated with *BRCA1* deficiency but also validate an approach, more applicable and convenient than previous strategies, to study break-induced TD formation.

## Methods

### Plasmids and chemical reagents

The pcDNA3β-Hyg-based plasmids expressing Cas9, Cas9^D^ and Cas9^H^ were previously generated (Feng *et al*, 2022). Plasmids expressing sgRNAs were constructed from the U6-sgRNA vector as described previously (Feng *et al*, 2022). The sgRNA target sequences are listed in Table EV 3. The TA cloning vector was purchased from TSINGKE (TSV-007S pClone007 Simple Vector Kit).The PARP inhibitor Olaparib was purchased from Selleck (S1060).

### Cell lines and cell culture

Mouse ES cells were grown in DMEM medium (Biological Industries) supplied with 20% fetal bovine serum (Gibco), 1% penicillin-streptomycin (Gibco), 2 mM L-glutamine (Gibco), 0.1mM β-mercaptoethanol (Sigma), 0.1mM non-essential amino acid (Gibco), 1mM sodium pyruvate (Gibco) and 1000U/mL leukemia inhibitory factor (Millipore) on either mouse embryonic fibroblast (MEF) feeders or gelatinized plates. Mouse ES cells containing the single-copy SCR-RFP reporter integrated at the *Rosa26* locus were previously established (Chandramouly *et al*, 2013; Rass *et al*, 2013). As described before (Feng *et al*, 2022; Nacson *et al*, 2020), isogenic mouse *Brca1^m/m^* and *Brca1^△cc/△cc^* ES clones were generated along with isogenic *Brca1^+/+^* clones *via* partial deletion of *Brca1* BRCT domain and CC domain by paired Cas9-sgRNA approach, respectively. Briefly, 2×10^5^ reporter mouse ES cells were transfected with the expression plasmids for Cas9 and paired sgRNAs targeting *Brca1* BRCT domain or CC domain in a 24-well plate. The target sequences of paired sgRNAs are listed in Table EV 3. At 72 h post transfection, 2,000 cells were seeded onto MEF feeders at a 10-cm plate for single clones without selection. After colony formation, single clones were picked and expanded, and gDNA was extracted with a purification kit (Axygen). Isogenic mouse *Brca1^+/+^*, *Brca1^m/m^*and *Brca1^△cc/△cc^* ES clones were confirmed by Sanger sequencing of targeted PCR amplicons with primers (Table EV 3) and by Western blot.

### Transfection and TD reporter assays

Transfection of mouse ES cells was performed with Lipofectamine 2000 (Invitrogen) in a 24-well plate as previously described (Xie *et al*, 2004; Willis & Scully, 2021). Total 2×10^5^ TD reporter mouse ES cells were transfected with 0.5 μg DNA. For the siRNA experiments, the same number of TD reporter mouse ES cells were transfected with 20 pmol siRNA together with 0.5μg expression plasmids for Cas9-sgRNA or Cas9n-sgRNA. Transfected or treated cells were analyzed for *RFP^+^* frequencies using the Beckman Coulter CytoFLEX flow cytometer at least 3 days post-transfection. Fluorescence-activated cell sorting (FACS) data were analyzed using the CytExpert 2.0 software. The TD frequencies were calculated by correcting *RFP^+^*frequencies with background readings and normalizing with transfection efficiencies as described before (Feng *et al*, 2022).

### Generation of TD reporter mouse ES cells

Mouse ES cells containing the single-copy SCR-RFP reporter integrated at the *Rosa26* locus were transfected with expression plasmids for paired Cas9-gRNAs which target sites flanking *TrGFP* in the SCR-RFP reporter. The target sequences of paired sgRNAs are listed in Table EV 3. After 72 h post-transfection, 2,000 cells were seeded on MEF feeder cells in a 10-cm plate for single clones without selection. After colony formation, single clones were picked and expanded, and gDNA was extracted with a purification kit (Axygen). The clone with precise deletion of *TrGFP* was determined by Sanger sequencing of targeted PCR amplicons with PCR primers (Table EV 3) and selected as the TD reporter clone.

### Western blot and immunofluorescence

For Western blot, cells were harvested after 72 h post-transfection and lysed with the RIPA buffer (20 mM Tris-HCl pH 7.5, 150 mM NaCl, 1 mM Na_2_EDTA, 1 mM EGTA, 1% NP-40, 1% sodium deoxycholate, 2.5 mM sodium pyrophosphate, 1 mM beta-glycerophosphate, 1 mM Na_3_VO_4_, 1 µg/mL leupeptin) for 30 min. Cell extractions were separated by SDS-PAGE electrophoresis and analyzed by Western blot. Primary antibodies used include anti-β-Actin (AT0001, 1:1000) from Engibody, anti-Rad51 (EPR4030, 1:1000) from Abcam, and anti-mouse Brca1 antibody(Wu *et al*, 2009).

For immunofluorescence staining, mouse ES cells were seeded on glass coverslips in a 6-well plate, cultured for 24 h, and treated with 5μM Olaparib as well as the mock control DMSO. After 6-h treatment, cells were fixed with 4% paraformaldehyde for 20 min, permeabilized with 0.1% Triton X-100 for 10 min, and blocked in 5% bovine serum albumin in PBS for 1 h. Cells were probed with primary antibodies including anti-Rad51 (EPR4030, 1:1000) from Abcam and anti-γH2AX (sc-517348, 1:200) from Santa Cruz and with Alexa Fluor 488-conjugated secondary antibodies (#111-545-003, 1:1000) or Alexa Fluor 594-conjugated secondary antibodies (#115-585-003, 1:1000) from Jackson ImmunoResearch, stained with 4′,6-diamidino-2-phenylindole (DAPI), and imaged by a fluorescence microscope (Leica DM4000).

### RNA interference and quantitative reverse transcription PCR

Small interference RNAs (siRNAs) targeting mouse *Brca1*, *Rad51*, *Brca2* and non-target siRNA (siNT) as a negative control (Table EV 3) were purchased from RiboBio Co. Total 2.0 × 10^5^ mouse ES cells were transfected with 20 pmol siRNA together with expression plasmids for Cas9-sgRNA or Cas9n-sgRNA. At 3 d post-transfection, RNAs were isolated and reverse-transcribed to complementary DNA using the HiScript II Q RT SuperMix for qPCR (Vazyme). Quantitative reverse transcriptase PCR (qRT-PCR) was performed for siRNA-mediated *Brca1* and *Brca2* depletion on qPCR CFX 96 Thermocycler (Bio-Rad) using gene-specific primers (Table EV 3). Depletion of *Rad51* was confirmed by Western blotting.

### Characterization of TD structures in individual RFP^+^ clones

At 72 h post transfection, TD reporter mouse ES cells were sorted for induced *RFP^+^* cells by FACS using Beckman Moflo Astrios EQ. Sorted cells were seeded on MEF feeder cells for single-clone isolation. gDNA were isolated from individual *RFP^+^* clones with FastPure Cell/Tissue DNA Isolation Mini Kit (Vazyme). The invasion point and the termination point of the TD product from these individual clones were amplified by PCR with an outward primer pair in the TD reporter (Table EV 3) and determined by Sanger sequencing. The sequences from individual *RFP^+^*clones and individual PCR product subcloned into the TA cloning vector were aligned to their respective reference sequences of the TD reporter and the Invasion type and the Termination type of TDs were thus assigned.

### PCR-based TD assays at the TD reporter and at a natural site

To perform PCR-based assays of Cas9-and Cas9n-induced TD at the TD reporter site and the three natural sites *H11*, *Rosa26* and *Col1a1*, mouse ES cells were transfected with expression plasmids for Cas9-sgRNA or Cas9n-sgRNA as indicated to induce site-specific two-ended DSBs or DNA nicks at the TD reporter and the natural sites, along with pcDNA3β-GFP used as transfection efficiencies. At 72 h post transfection, cells were harvested and gDNA isolated from these cells with FastPure Cell/Tissue DNA Isolation Mini Kit (Vazyme). The points for invasion or termination in TD product were amplified by PCR with 50 ng gDNA and an outward primer pair, 10 μM each primer, in a 30-μL reaction. The outward primer pair is complementary to the sites upstream and downstream of Cas9 or Cas9n target sites, respectively. After 72 h, cells were harvested and gDNA was isolated for PCR. After PCR products were separated by agarose gel electrophoresis, the intensity of DNA bands in each lane was quantified by ImageJ and subtracted by background intensity. The *Gapdh* region was also amplified by PCR with the same amount of gDNA in the same condition of PCR reaction except different primer pairs as the internal control. The TD intensity relative to *Gapdh* was calculated as the ratios of the TD intensity to the *Gapdh* intensity.

To determine the TD structure in the TD reporter, DNA bands were excised and subcloned into a TA cloning vector after PCR products were separated by agarose gel electrophoresis. To determine the structure of TDs at the natural sites, PCR products were purified by the PCR clean-up kit (AxyPrep) and subcloned into a TA cloning vector (TSINGKE TSV-007S pClone007 Simple Vector Kit). Individual PCR products subcloned into the TA cloning vector were analyzed by Sanger sequencing. The sequences were aligned to the reference sequences of the TD reporter and the natural sites by the BLAST alignment tool and the Invasion type and the Termination type of TDs were thus assigned.

### Determination of TD span size

In each TD event, the TD junction identified by Sanger sequencing at or near the Cas9 or Cas9n target site revealed the invasion point for TD-inducing non-allelic DNA synthesis and was considered as the starting point of TD. The distance from this starting point to the next starting point of identical sequence was estimated to predict the length of DNA duplicated. TD span sizes were derived by doubling the predicted length of DNA duplicated. To determine the distribution of TD span sizes, the percentage of the number of TDs with a specific TD span size to the number of total TD events was calculated, and the density in a violin plot for a specific TD span size is derived by 100 x the percentage of the number of TDs with this specific TD span size to the number of total TD events.

## Data availability

Flow cytometry raw data and microscope datasets has been deposited at the Zenodo accessible at https://doi.org/10.5281/zenodo.10252147. Source data are provided with this paper.

## Acknowledgements

Anti-mouse Brca1 antibody is a generous gift from the Linyu Lu lab (Zhejiang University School of Medicine). We are grateful to helpful discussion with members of the Xie lab. We thank Bi Chao and Hong Xiaoli from the Core Facilities, Zhejiang University School of Medicine for their technical support. This work is funded by the National Natural Science Foundation of China (No. 32371348 to A.-Y.X. and No. 32071439 to Y.-L.F.) and the Department of Science and Technology of Hangzhou (202204A05 to A.Y.X. and 202204B08 to Y.L.F.).

## Declarations

### Ethics approval and consent to participate

Not applicable

### Consent for publication

Not applicable

## Authors’ contributions

A.-Y.X. conceived the project and supervised the study. Z.-C.H., Y.-L.F. and Q.L. generated DNA constructs and cell lines, and Z.-C.H. performed experiments and statistical analysis. R.-D.C., S.-C.L. and M.W. assisted with generation of DNA constructs and cell lines. A.-Y.X., Z.-C.H. and Y-L.F. analyzed the data. Z.-C.H. and A.-Y.X. made figures and tables, and A.-Y.X. and Z.-C.H. wrote the manuscript.

